# Viral Syncytia Evolve to Resist Interferon

**DOI:** 10.1101/2023.12.12.571262

**Authors:** Tiansheng Li, Insung Kang, Juan Ye, Zhe Hu, James Gibbs, Chengjin Ye, Kazuyo Takeda, Ivan Kosik, Guoli Shi, Jaroslav Holly, Martina Kosikova, Zhiping Ye, Alex A. Compton, Luis Martinez-Sobrido, Reed F. Johnson, Hang Xie, Jonathan W. Yewdell

## Abstract

SARS-CoV-2, like many viruses, generates syncytia but the role of syncytia formation in viral evolution remains unknown. Using SARS-CoV-2 and SARS-CoV-2 Spike (S) replacement vesicular stomatitis (VSV), we show that S-mediated syncytia impair the antiviral effects of interferons in cultured cells, human lung cell cultures, and hACE2 transgenic mice. Amino acid substitutions that modulate syncytia formation in Delta- and Omicron-encoded S have parallel effects on viral interferon resistance. S-mediated syncytia compromise antibody-mediated virus neutralization in cultured cells. We recapitulate interferon and neutralizing antibody resistance in syncytia generated by the orthoreovirus p14 fusion-associated small transmembrane (FAST) protein in VSV, influenza virus, and seasonal coronavirus OC43 infections. These findings explain selection of SARS-CoV-2 fusogenic variants in humans and, more generally, the evolution of fusogenic viruses driven by adaptive and innate immunity.

## INTRODUCTION

Many medically important enveloped and non-enveloped viruses induce fusion of infected cells with surrounding cells to generate syncytia: multinucleated giant cells^1^. Though syncytia-forming viruses are frequently selected by propagation in patients or cell lines, this is often paradoxically associated with decreased production of infectious viral progeny^2–5^. In this study, we examine this paradox as exhibited by SARS-CoV-2 and other viruses.

While circulating in billions of humans, SARS-CoV-2 continuously evolves immune evasion variants^6–11^. Much of the variation occurs in the Spike (S) virion surface glycoprotein, which facilitates the attachment of the virus to cells and subsequent fusion of viral and cell membranes. S possesses an RXXR furin cleavage site (FCS), whose cleavage generates the S1 and S2 subunits (**Fig. 1a**). This feature distinguishes SARS-CoV-2 from most related viruses in the Betacoronavirus family, although similar FCSs are present in more distantly related coronaviruses (**Fig. 1a**)^12,13^. FCS cleavage by furin and/or TMPRSS2 enhances viral entry by promoting fusion with the host cell membrane and enhances syncytia formation in cell cultures and in COVID-19 patients^14–19^. The FCS is strongly selected during SARS-CoV-2 circulation in humans (**Fig. 1b** and **Supplementary Fig. 1**) and is associated with increased pathology^20–25^. Since syncytia formation typically reduces viral replication in cultured cells, the basis for its positive selection in viral evolution is an important question.

**Fig. 1.**
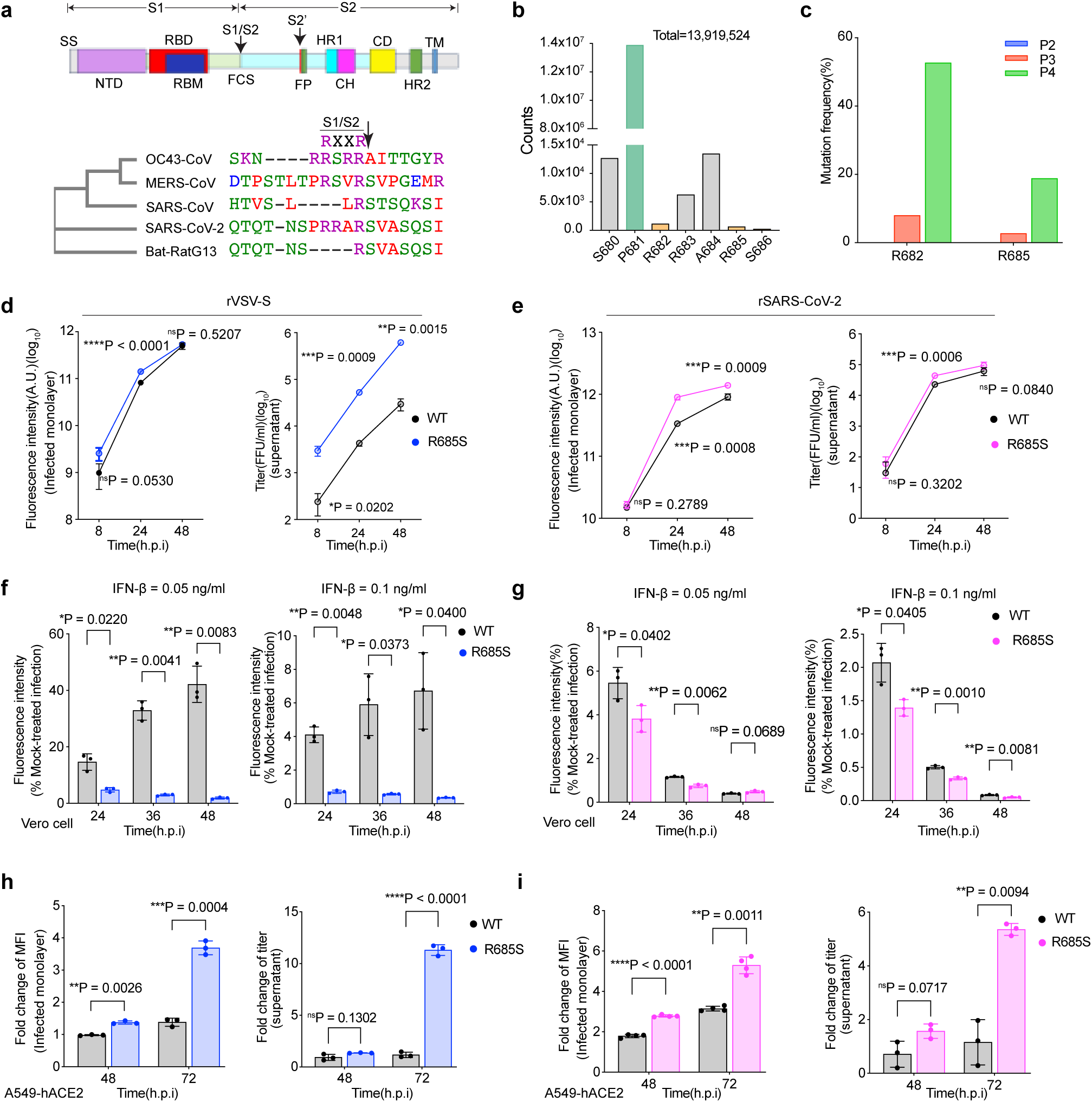
Furin cleavage site confers VSV and SARS-CoV-2 resistance to IFN-β anti-viral activity in cell culture. **a** Schematic of SARS-CoV-2 S protein structure(up, created by Adobe Illustrator), signal sequence (SS); N-terminal domain (NTD); receptor-binding domain(RBD); receptor-binding motif(RBM); S1/S2 protease cleavage site/furin cleavage site(S1/S2 FCS); S2’ protease cleavage site(S2’); fusion peptide(FP); heptad repeat 1(HR1); central helix(CH); connector domain(CD); heptad repeat 2(HR2); transmembrane domain(TM). Polygenetic tree and alignment of S FCS region sequences from five β coronaviruses (bottom). The conserved two key residues RXXR in FCS and cleaved residue are arrow-headed. **b** Mutations at FCS in patient SARS-CoV-2 isolates. Substitution counts at each residue at or near the FCS SARS-CoV-2 genome sequences deposited to the GISAID database as of May 26^th^, 2023. **c** Mutation frequency at FCS in passaged recombinant replication-competent rVSV-S virus in BHK21-ACE2 cells. The viral genomes of passage 2 to 4 viral stocks were subjected to next-generation sequencing to identify mutations in the *S* gene, revealing mutations in only FCS. **d** and **e** Vero cells were infected by rVSV-S (**d**) (n=3 biological replicates) or rSARS-CoV-2(**e**) (n=3, or 4 biological replicates for FI and titer, respectively). Data show mean ± s.d. **f** and **g** Sensitivity of *wt* and R685S -bearing viruses to exogenous IFN-β in Vero cells. Cells were treated with IFN-β for 20 h prior to infection by rVSV-S (**f**)(n=3 biological replicates) or rSARS-CoV-2(**g**)(n=3 biological replicates) at a MOI of 0.01. Data show mean ± s.d. **h** and **i** Blocking JAK1/2 with Ruxolitilib (ruxo) enhances R685S replication in A549-ACE2 cells. A549-ACE2 cells were treated with 2 μM ruxo for 2 h prior to infection by rVSV-S(**h**) or rSARS-CoV-2(**i**) at MOI of 0.01, with ruxo maintained throughout the infection. Fold change of mean fluorescence intensity (MFI) of infected cells (left) and fold change of titer in the supernatant (right) were normalized by mock-treated values. n=3 biological replicates, except the left panel of **i**. in which n=4 biological replicates per group. Data show mean ± s.d. Statistical analysis was performed using a two-tailed, unpaired t-test with Welch’s correction. *P < 0.05, **P < 0.01, ***P < 0.001 and ****P < 0.0001; ns, not significant.

Interferons play a central role in limiting viral replication in the initial stages of viral infection, particularly in naïve individuals. Strong evidence exists that interferons are a major factor in controlling SARS-CoV-2 ^26–28^ and other human viruses ^29^. Studying the evolutionary selection of the FCS *in vitro* and *in vivo*, we have found a critical role for syncytia formation in evading the anti-viral activity of interferons (IFNs) and Abs (Abs).

## RESULTS

### FCS impairs virus replication

To better understand the contribution of the S FCS to viral fitness, we replaced the receptor gene of vesicular stomatitis Indiana virus (VSV) with the ancestral SARS-CoV-2 *S* gene to generate a replication-competent VSV-eGFP-SARS-CoV-2 S virus (referred to as rVSV-S, **Supplementary Fig. 2a**). We removed the coding sequence for the S 21-residue C-terminal ER/GC retention sequence to increase S cell surface expression and incorporation into VSV virions^30,31^. We passaged the initial virus stock (P0) in BHK21 cells expressing human ACE2 (referred to as BHK21-ACE2) for several passages. Sanger sequencing at passage 2 (P2) showed only *wt* (Wuhan-Hu-1 spike) virus. We detected a small population of the mutant G2045A, corresponding to an R to Q substitution at position 682 (R682Q) after passage 3 (P3), becoming dominant in P4 and P5, accompanied by G2054A, corresponding to R685H (**Supplementary Fig. 2b**).

We next sequenced the *S* genes of 22 plaque-purified P4 stocks expanded in BHK21-ACE2 or MA104 cells, revealing four FCS loss mutants, R685S, R682W, R685H, and R682Q. Each mutant attained higher titers in BHK21-ACE2 cells than the *wt* virus (**Supplementary Fig. 2c**). Next-generation sequencing (NGS) revealed an increasing mutation frequency with passage number in BHK21-ACE2 cells occurring at bases encoding R682 or R685 (**Fig. 1c**). Extending previous findings of FCS loss during propagation in Vero cells for both rVSV-S and rSARS-CoV-2^22,30–32^, this indicates that FCS substitutions enhance virus replication in BHK21-ACE2 cells. To explore the context-dependent fitness of the FCS, we focused on the R685S mutant among the four selected mutants, as they exhibited similarly reduced syncytia formation (**Supplementary Fig. 2c**). Western blot analysis of rVSV-S-infected Vero cells revealed that the R685S mutation resulted in an almost complete loss of cleavage at the FCS (**Supplementary Fig. 2d**). A virus growth competition assay using an increasing mutant to *wt* virus ratio demonstrates positive selection for R685S *vs. wt* virus in BHK21-ACE2, and Vero cells and negative selection in Calu-3 and Caco-2 cells (**Supplementary Fig. 2e**).

To minimize the effects of possible passenger mutations, we generated stocks of recombinant rVSV and rSARS-CoV-2 fluorescent protein (FP) reporter viruses expressing *wt* or R685S S, confirming viral genome sequence by NGS. After low multiplicity of infection (MOI of 0.01) of Vero cells, R685S-virus replicated faster than *wt* virus in both rVSV and rSARS-CoV-2 systems (**Fig. 1d** and **1e**) as determined by released virions and microscope-based quantitation of reporter FP expression in either live (rVSV-S) or paraformaldehyde (PFA) fixed cells (for rSARS-CoV-2). Importantly, imaging also showed that the size of R685S infection foci was significantly greater than *wt* S in both SARS-CoV-2 or VSV-S vectors (**Supplementary Fig. 3a to f**).

### FCS fitness is IFN-dependent

We noted that FCS loss is selected by virus passaging in Vero and BHK21 cell lines, which share a deficiency in IFN-secretion^33,34^. We tested the contribution of IFN to FCS evolution by adding IFN-β to Vero cells and infecting the cells 20 h later with *wt* or R685S rVSV-S or rSARS-CoV-2 viruses (**Fig. 1f** and **1g**). As expected, rVSV-S and rSARS-CoV-2 were sensitive to IFN-β treatment, exhibiting at least 80% inhibition of FP-reported infection 24 h post-infection (h.p.i) at the lower IFN-β dose used and >96% inhibition at the higher dose. Importantly, relative to R685S, *wt* S conferred up to 23-fold resistance to IFN-β in rVSV-S and less (up to 1.5-fold), but statistically significant resistance in rSARS-CoV-2.

Extending these findings to an IFN secretion-competent cell (A549-ACE2), we used the JAK1/2 kinase small molecule inhibitor Ruxolitinib to block IFN-mediated signaling. For both *wt* rVSV-S and rSARS-CoV-2 infections, measuring either infected cell FP signal or released infectious virus, Ruxolitinib enhanced R685S infection to a much greater extent than *wt* infection (**Fig. 1h** and **1i**).

Taken together, these data support the conclusion that the S FCS confers IFN-β-resistance to VSV and SARS-CoV-2 in cultured cells.

### Syncytia evade IFN-induced anti-viral activity

How does the FCS confer resistance to IFN-β anti-viral activity? After 18-20 h IFN treatment, we infected cells with rVSV-S and determined the half maximal inhibitory concentration (IC_50_) of IFN-β in primary infected cells by *in situ* imaging before the release of virus particles (7 h.p.i) (*wt*, 1.16 pM; R685S, 1.14 pM) (**Fig. 2a**). Compared with rVSV-S, rSARS-CoV-2 was slightly less sensitive to IFN-β, but *wt* and R685S exhibited near identical IC_50_ values (2.39 *vs.* 2.13 pM) (**Fig. 2b**). These findings indicate that the FCS does not affect IFN-β inhibition of initial viral entry in Vero cells.

**Fig. 2.**
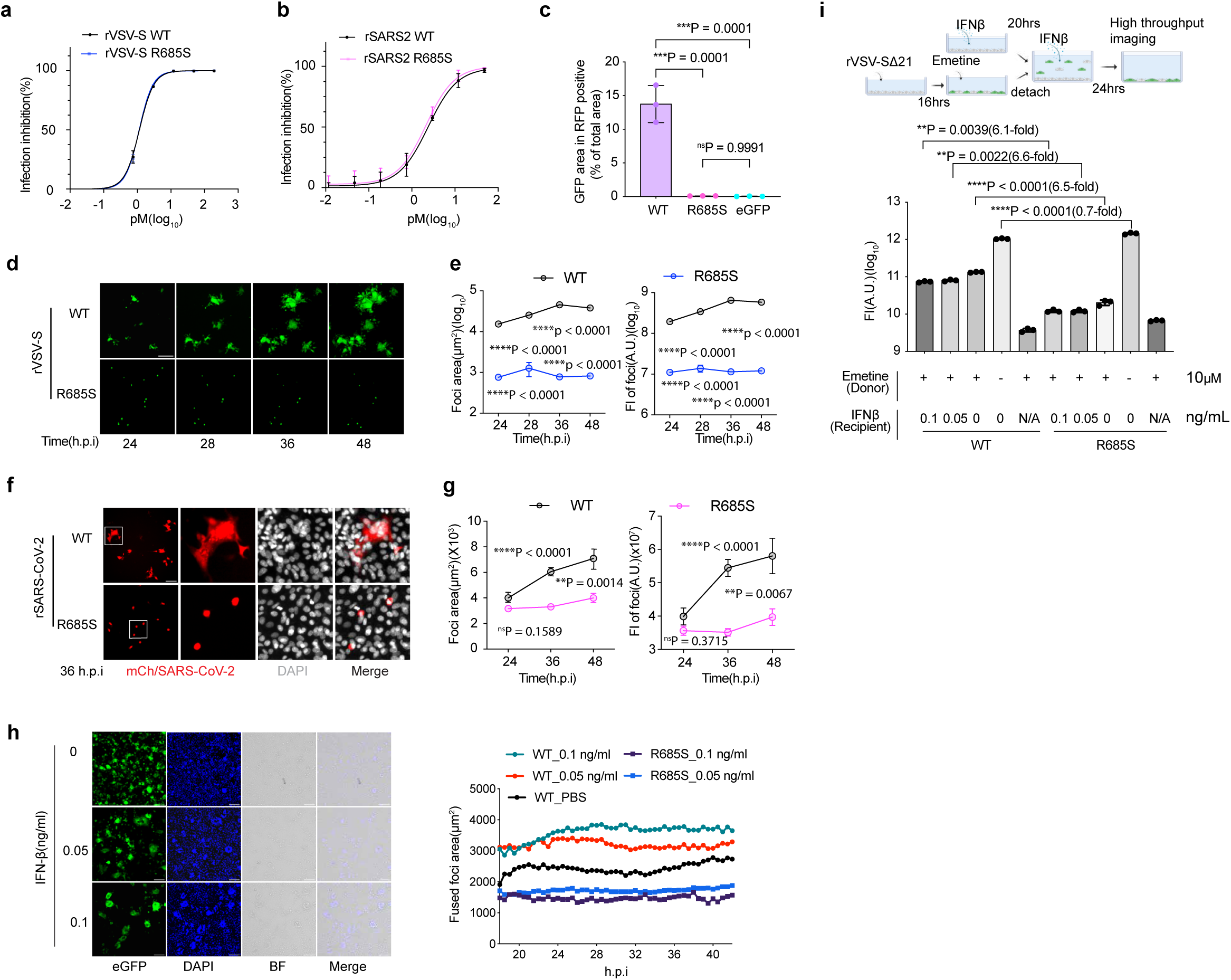
Syncytia confer virus resistance to IFN-β anti-viral activity. **a** and **b** Effect of exogenous IFN-β to viral entry on *wt* and R685S in rVSV-S (**a**) and rSARS-CoV-2 (**b**) was measured in Vero cells at 7 h.p.i with a MOI of 0.01. Series diluted IFN-β was added to monolayers 20 h prior to inoculation. Data show mean ± s.d (n=3 biological replicates). **c** Syncytia mediated by S in Vero cells. Syncytial area as determined by GFP-RFP overlapping pixels. Data show mean ± s.d (n = 3 biological replicates). **d** Time-lapse imaging of rVSV-S infection in Vero cells exposed to IFN-β (0.025 ng/ml) 20 h prior to infection, at MOI of 0.04. Scale bar, 250 μm **e** The area (left) and fluorescence intensity (right) of infection foci in **d**. Data show mean ± s.e.m (n = 500 foci). **f** Representative images of rSARS-CoV-2 *wt* and R685S infections in Vero cells under IFN-β treatment (0.1 ng/ml) at 36 h.p.i. Higher-magnification views of the regions indicated by squares are shown in the right images. Scale bar, 100 μm **g** Area (left) and fluorescence intensity (right) of infection foci in **f**. Data show mean ± s.e.m (n = 128, 163, 88 foci for *wt*; 165, 156, 78 foci for R685S, at 24, 36, and 48 h.p.i, respectively). **h** Syncytia formation of rVSV-S *wt* infection in Vero cells under IFN-β treatment. Representative images at 48 h.p.i. (up), and time-lapse measure of mean area of fused focus (setting: object size>60 μm in Gen5) from 18 h.p.i to 48 h.p.i(bottom) (n = 24 fields per group). **i** rVSV-S bypasses emetine-blocked translation via syncytia formation. Experimental design is summarized at the top, with details in Materials and Methods. Schematic diagram was created in BioRender (Gibbs, J. (2026) https://BioRender.com/wpfype0). Data show mean ± s.d (n = 3 biological replicates). N/A, no receptor cell. FI, florescence intensity. A.U., arbitrary units. Statistical analysis was performed using a one-way ANOVA (**c**) or two-way ANOVA (**e**, **g** and **i**) with multiple comparisons. **P < 0.01, ***P < 0.001 and ****P < 0.0001; ns, not significant.

Rather, we found that the FCS reduces IFN anti-viral activity by favoring syncytia formation. We inferred this initially by overlaying S +EGFP transfected Vero cells with RFP transfected Vero cells. Measuring cell fusion by imaging revealed that maximal cell fusion requires S with a functional FCS (**Fig. 2c**), extending prior reports ^30,31^. To further establish the role of the FCS-mediated fusion in evading IFN, we live-imaged infected Vero cells to measure rVSV-S dissemination. When we treated cells with IFN-β, *wt* rVSV-S infectious foci enlarged over time, while rVSV-S R685S foci remained limited to single infected cells, many of which died during imaging (**Fig. 2d** and **2e** and **Supplementary Fig. 4**). With rSARS-CoV-2 wt infection using 0.1 ng/ml IFN-β treatment (**Fig. 2f** and **2g** and **Supplementary Fig. 5**), we observed much larger infection foci consisting of highly multinucleated cells with higher fluorescence relative to R685S infection. These data demonstrate that IFN-β under the tested concentration does not block S-mediated Vero cell syncytia formation.

Comparing IFN-β *vs.* control treatment in rVSV-S infection imaging experiments (**Supplementary Fig. 6a**), we observed that IFN-β blocks dissemination to non-adjacent cells, consistent with its entry blockade in experiments (**Fig. 2a**). Treating Vero cells with IFN-β increased the fusion ratio (defined as the area of syncytia divided by the area of syncytia + single infected cells) 1.5-fold vs untreated cells (90% *vs* 60%) at 24 h.p.i. (**Supplementary Fig. 6a**). Overlaying cells with agar to prevent diffusion of released virus did not affect the fusion ratio of IFN-β treated cells while reducing the fusion ratio of untreated cells to 35% (**Supplementary Fig. 6a**). The IFN-β-mediated fusion ratio increase predominantly reflected increases in the size of individual syncytia (containing more individual cells) rather than the number of syncytia (**Fig. 2h**, and **Supplementary Movie 1 to 4**). In contrast to rVSV-S *wt*, rVSV-S R685S virus exhibited few and small syncytia, with or without IFN-β (**Fig. 2h** and **Supplementary Fig. 6b**).

To explore the contribution of donor (virus-infected) *vs.* recipient cell protein synthesis to virus replication following syncytia formation we used emetine to irreversibly block protein synthesis^35,36^ in donor Vero cells 8 h post-infection with rVSV-S. After removing emetine, we overlaid infected cells on uninfected Vero cells in media with decreasing amounts of IFN-β (**Fig. 2i**). After 24 h, we measured GFP expression in cultures by high throughput imaging. As expected, donor cells expressing *wt* S fused with receptor cells to form multinucleated syncytia with or without IFN-β at much higher levels than donor cells expressing R685S S (**Supplementary Fig. 6c**). Treating donor cells with emetine did not affect the large discrepancy between *wt* and R685S in generating syncytia, as determined by image analysis quantitation of syncytia per field or the area of GFP expressing foci (**Supplementary Fig. 6d**). Adding emetine-treated *wt* S infected cells to uninfected cells increased overall GFP expression 36-fold in the absence of IFN, a value that was only slightly reduced (1.7-fold) by maximal IFN treatment. By contrast, adding R685S S emetine-treated infected cells to uninfected cells only increased total GFP expression by less than 2-fold with or without IFN, attributable to limited release of infectious virions. Thus, greater than 90% of the GFP synthesized by *wt* virus in syncytia derives from recipient cell ribosomes.

Together, these results demonstrate that:

1. IFN-β selectively inhibits S-mediated cell-free *vs.* syncytia-based transmission of SARS-CoV-2 and VSV.
2. Syncytia rapidly amplify viral gene expression by increasing the number of ribosomes capable of translating viral mRNAs.

### Syncytia confer SARS-CoV-2 IFN resistance in human primary small airway epithelial cells

We extended our findings to human primary small airway epithelial cells maintained in an air-liquid interface (ALI), a more biologically relevant system. Since airway epithelia have receptors for type III IFNs, which play an important role in respiratory infections^37^, we included IFN-λ2 in experiments. Four days after infecting cells with rSARS-CoV-2, we imaged virus-encoded mCherry as a measure of viral gene expression and determined titers of released virus. In the absence of IFNs, *wt* and R685S virus replicated nearly identically (**Fig. 3a**). Exposing cells before and after infection to either IFN-β or IFN-λ2 reduced *wt* rSARS-CoV-2 replication 2- and 6-fold, respectively, by either criterion. Importantly, both IFNs had a much greater effect on blocking R685S rSARS-CoV-2 replication (**Fig. 3b** and **3c**). Imaging infected cells confirmed syncytia formation by *wt* but not the R685S virus (**Fig. 3d**).

**Fig. 3.**
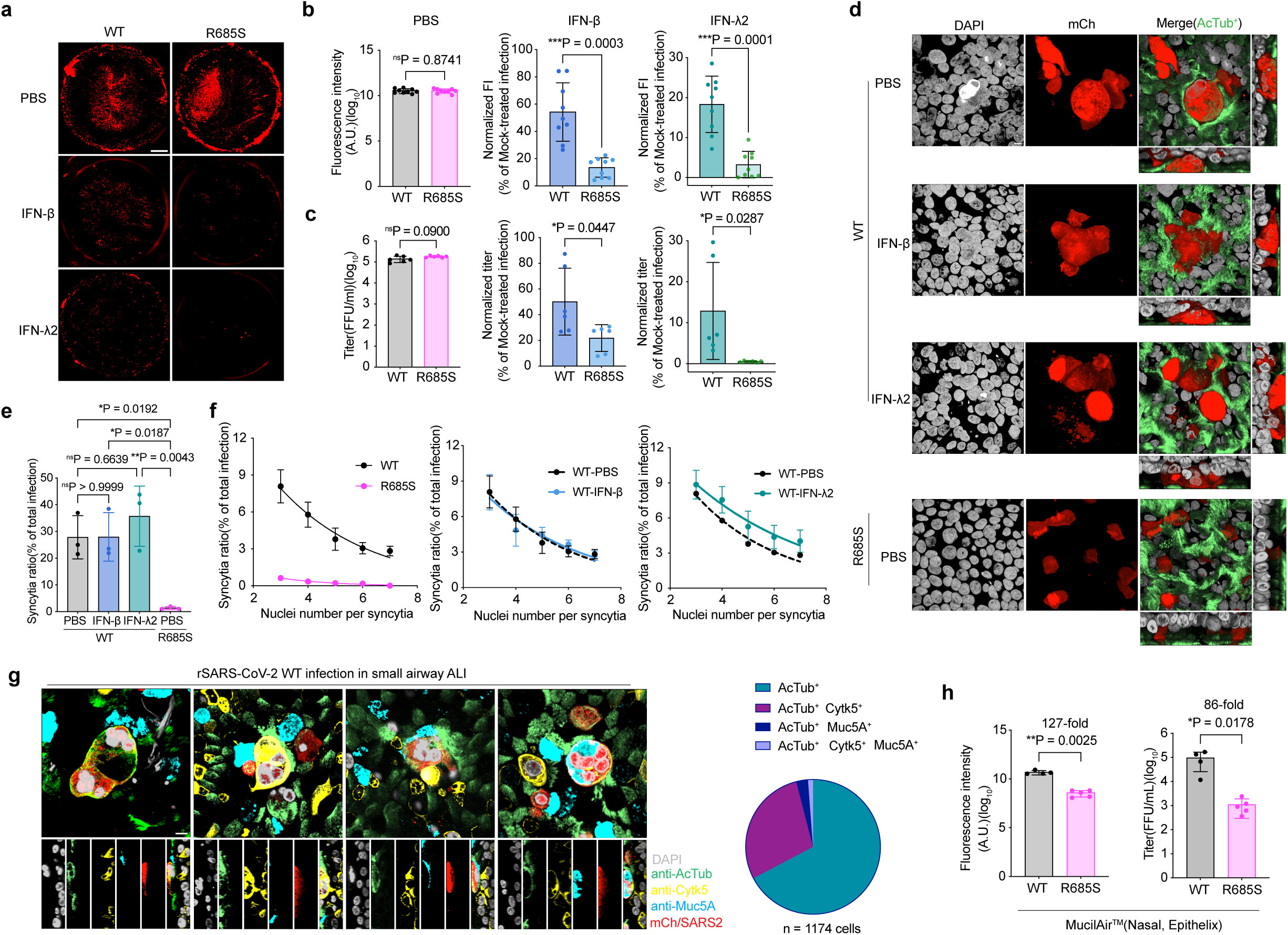
FCS confers SARS-CoV-2 resistance to IFNs in human small airway epithelial cells and in hACE2 transgenic mice. **a-c** Sensitivity of rSARS-CoV-2 *wt* and R685S to IFNs in small airway epithelial cell in air-liquid interface (ALI) format. Representative images of infection in entire ALI cultures treated with IFN-β (0.2 ng/ml) or IFN-λ2 (10 ng/ml) at 4 d.p.i (**a**). Scale bar, 1000μm. Fluorescence Intensity of infected cells treated with PBS (left), IFN-β(middle) or INF-α2 (right) (**b**, n= 9 tissues). Titer of apical wash from infected cells treated with PBS (left), IFN-β (middle) or IFN-λ2 (right) **(c**, n=6 tissues). 3 donors for each group. Data show mean ± s.d. **d** Representative images of syncytia formation in infected small airway ALI treated with IFN-β or IFN-α2 at 4 d.p.i. ALI was stained with anti-AcTub (ciliated cells, green signal) and counterstained with DAPI (gray signal). rSARS-CoV-2 was shown by red signal. Z projection was shown. Scale bar, 5μm. **e** Syncytia ratio of rSARS-CoV-2 infection in small airway ALI with IFN treatment at 4 d.p.i. The number of individual infected and syncytial cell nuclei was determined from reconstructed 3D confocal images (syncytia defined as 3 or more nuclei in one infected focus). 28 to 47 fields per group were analyzed. Data show mean ± s.d (n=3 tissues). **f** The nuclei number per syncytia in rSARS-CoV-2 infected in small airway ALI. Data were fit in Gaussian distribution and shown in mean ± s.d (n=3 tissues). **g** Cell types present in syncytia. Infected small airway ALI were stained with anti-AcTub (green signal, marker for ciliated cell), anti-Cytk5 (yellow signal, marker for basal cell) and anti-Muc5A (cyan signal, marker for goblet cell) Abs and counterstained with DAPI (gray signal). The distribution of cell types in syncytia is shown by the pie chart. Scale bar, 5μm. **h** Comparison of rSARS-CoV-2 *wt* and R685S replication in primary nasal epithelial cells in ALI format (MucilAir from Epithelix) at 4 d.p.i. Data show mean ± s.d (n=4 tissues). Statistical analysis was performed using a two-tailed, unpaired t-test (**b,c,h**) or one-way ANOVA (**e**) with multiple comparisons.

We quantitated the effects of IFN on rSARS-CoV-2-induced syncytia using image analysis software. No differences in syncytia ratio were observed after IFN-β treatment at a concentration of 0.2 ng/ml, while a slight increase in syncytia ratio occurred in the IFN-λ2 group compared to the group without IFN treatment. Importantly, rSARS-CoV-2 R685S rarely generated syncytia without IFN (**Fig. 3e**). The nuclei count in each syncytium of *wt* ranged from 3 to more than 8, fitting a Gaussian distribution. IFN-β treatment at a concentration of 0.2 ng/ml did not alter the distribution of nuclei in syncytia. In comparison, IFN-λ2 at a concentration of 10 ng/ml slightly increased the overall syncytia ratio (**Fig. 3f**), likely because of high dose of IFN-λ2 strongly inhibits cell-free infection and thus increases the syncytia ratio. Overall, this suggests that IFN selectively inhibits non-syncytial infection, favoring syncytia in human small airway epithelial cells, consistent with our results in Vero cells and a previous study^38^.

To determine the cell types that are involved in syncytia driven by SARS-CoV-2 infection, we stained for markers of ciliated cells (AcTub), goblet cells (Muc5AC), and basal cells (Cytk5). This revealed that syncytia form between ciliated cells, the primary target for SARS-CoV-2 (67%), ciliated cells and basal cells (29%), ciliated cells and goblet cells (3%), or between all three cell types (1%) (**Fig. 3g** and **Supplementary Fig. 7a**).

We next examined SARS-CoV-2 replication in nasal epithelial ALI cultures. Interestingly, while *wt* rSARS-CoV-2 replicated to a similar extent as in small airway ALI cultures (determined by fluorescence intensity and infectious virus released 4 d.p.i, **Fig. 3h** and **Supplementary Fig. 7b**), R685S rSARS-CoV-2 virus infectious titers were 86-fold lower in nasal cultures, indicating that R685S replication is attenuated in upper airway epithelial cells, suggesting a more vigorous IFN response in the upper *vs.* lower airway which is in line with previous studies^39^.

### *In vivo* evidence for SARS-CoV-2 syncytia as a viral IFN-escape strategy

Extending these findings *in vivo*, we intranasally infected K18-hACE2 transgenic mice using conditions leading to viral pneumonia. To examine the effects of IFN, we treated mice with IFN-λ2, chosen based on preferential expression of type III IFNs in virus-infected epithelial cells, milder pro-inflammatory effects *vs.* type I and II IFNs (confirmed in **Supplementary Fig. 8a**), and published efficacy in the mouse SARS-CoV-2 model^40^. Consistent with the small airway ALI culture experiments, *wt* and R685S rSARS-CoV-2 viruses replicated similarly in IFN-untreated mice, as determined by vRNA/infectious virus recovered from the lung (**Fig. 4a, b**). Intranasal IFN-λ2 treatment on day −1 and +1 d.p.i. had a greater effect on R685S *vs. wt* replication, with a 2-fold greater reduction in vRNA and 9-fold greater reduction in infectious virus (**Fig. 4a, b**). *Wt* rSARS-CoV-2 and R685S infection induced similar pulmonary proinflammatory cytokine expression profiles 3 d.p.i., except for an increase in *wt* virus-induced IL-5 and IFN-γ (**Supplementary Fig. 8b**). For both viruses, IFN-λ2 treatment decreased proinflammatory cytokines, consistent with reducing virus replication.

**Fig. 4.**
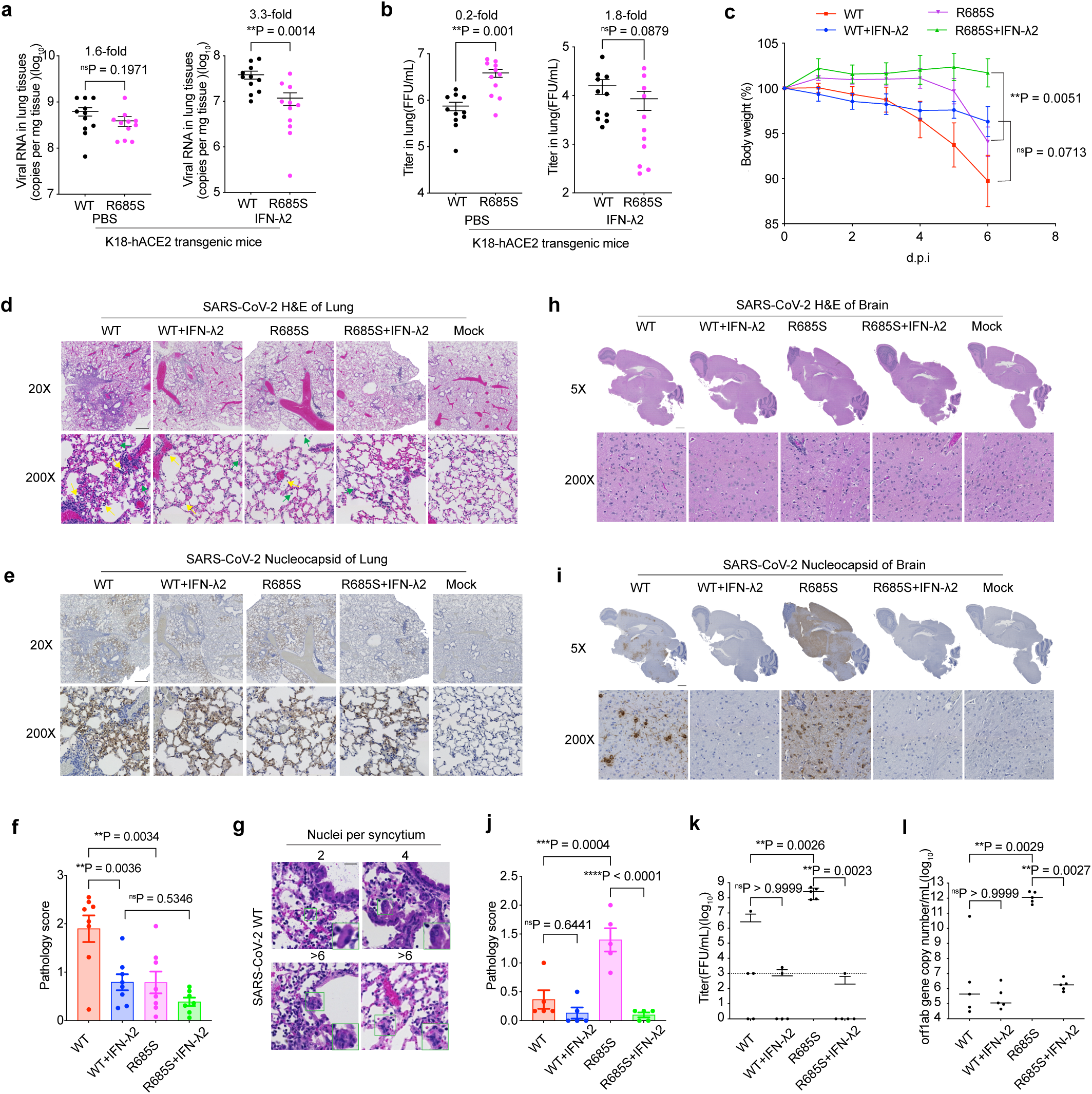
Histopathological analysis of IFN-λ2-mediated protection against wt and R685S SARS-CoV-2 infection in K18-hACE2 mice. **a-b** Viral load of rSARS-CoV-2 *wt* and R685S infection in lung tissue of K18-hACE2 mice with two doses of IFN-α2 or PBS treatment. Data show mean ± s.e.m (n=11 mice). **c** K18-hACE2 mice were intranasally infected with rSARS-CoV-2 WT or R685S and treated with IFN-λ2 or PBS. Body weight was monitored daily for 6 days. Data represent mean ± s.e.m. (n = 8 mice per group, except R685S + IFN-λ2 group, where n = 7 mice) **d and h** Hematoxylin and eosin (H&E) staining of lung (**d**) and brain (**h**) sections from K18-hACE2 mice at 6 d.p.i. with rSARS-CoV-2 WT, R685S, or mock infection. Representative images are shown at low magnification (top; 20×, scale bars: 500 μm; 5×, scale bar:1mm) and high magnification (bottom; 200×). Lymphocyte infiltration (yellow arrows) and mononuclear cell infiltration (green arrows) are evident in infected tissues. **e and i**, Immunohistochemical staining for SARS-CoV-2 nucleocapsid protein in lung (**e**) and brain (**i**) sections from infected or mock-infected K18-hACE2 mice at 6 d.p.i. Representative images are shown at low magnification (top panels; 20×, scale bars = 500 μm, 5×, scale bar = 1mm) and high magnification (bottom panels; 200×). **f and j** Pathology evaluation of infected mice lung (**f**) (n = 8 mice per group, except R685S + IFN-λ2 group, where n = 7 mice) and brain (**j**) (n = 5 mice). **g** Syncytia formation (highlighted by a green frame in the inset at higher magnification) was observed in lung sections of K18-hACE2 mice infected with SARS-CoV-2 *wt* at 6 d.p.i. The number of nuclei within each syncytium varied as indicated. Scare bar = 100 μm. **k and l** Viral loads of rSARS-CoV-2 *wt* and R685S in brain tissue of K18-hACE2 mice treated with IFN-λ2 or PBS. Infectious viral titers (**k**) and viral RNA copies (**i**) are shown. Data show mean ± s.d (n=5 mice). Statistical analysis was performed using a two-tailed, unpaired t-test (**a,b**), two- (**c**) or one- (**f,j,k,l**) way ANOVA with multiple comparisons. Results represent pooled data from two independent experiments.

To evaluate pathogenicity and syncytia formation, we infected K18-hACE2 transgenic mice with 10,000 FFUs of wt or R685S virus, giving a single i.n. dose of IFN-λ2 or PBS at 4 h.p.i. After monitoring weight loss for six days, we euthanized mice and performed histopathological analysis of lung and brain. *wt* virus was more pathogenic in the lungs than the R685S mutant, as indicated by greater body weight loss, increased immune cell infiltration, and higher viral nucleocapsid abundance (brown staining, blue is hematoxylin-stained nuclei)(**Fig. 4 c-f**). IFN-λ2 treatment provided protection against disease for both viruses but with a significantly stronger effect on the R685S mutant, leading to reduced weight loss, decreased lung pathology, and lower viral replication (**Fig. 4 c-f**). Histological analysis of lung tissue revealed occasional syncytia formation in the *wt*-infected group, with syncytia containing varying numbers of nuclei (**Fig. 4g**). In brain tissue, viral replication was detected in only 1 out of 5 mice infected with the *wt* virus, as shown by the presence of infectious viral progeny and positive N protein staining. In contrast, all 5 mice infected with the R685S mutant exhibited robust viral replication and pathogenicity in both the cerebrum and brainstem(**Fig. 4 h-l**). Furthermore, we observed spongiosis accompanied by neuronal degeneration and necrosis in the cerebral cortex of R685S-infected mice (**Supplementary Fig. 8c**).

These findings indicate that while the fusogenic *wt* virus evades IFN and replicates more efficiently, causing greater pathology in the lungs, the FCS mutant R685S exhibits enhanced neurotropism and brain pathogenicity, potentially due to lower endogenous interferon responses in the central nervous system in conjunction with its increased replication in the absence of IFNs^41^.

Together, these findings support the conclusion that the FCS improves viral fitness *in vivo* by enhancing escape from IFN, presumably due to enhancing syncytia formation.

### Syncytia enhance viral resistance to antibody neutralization

To investigate whether syncytia alters anti-S antibody viral infectivity neutralization (VN), we devised an imaging-based assay that uniquely records VN in real-time. We used Vero-ACE2 cells for rVSV-S infection and human lung alveolar basal epithelial A549/ACE2/TMPRSS2 (ACE2plusC3) cells for rSARS-CoV-2 infection. Both cell lines exhibit enhanced syncytia formation due to transgene-encoded protease overexpression. We tested six potent monoclonal Abs (mAbs) specific for the S receptor binding domain (RBD)^42,43^ for their ability to block virus-encoded reporter eGFP or mCherry expression. Abs were mixed with virus before infection and maintained during 20 h infection. Data are expressed as the Ab neutralization dose (ND_50_) required to reduce the FP signal by 50% relative to the no Ab control. We collected images at 7 and 20 h.p.i. (**Supplementary Fig. 9**)

Using rVSV-S to infect Vero-ACE2 cells, large syncytia formed at 20 h.p.i for *wt* virus but not for R685S virus (**Supplementary Fig. 9a**). Abs demonstrated negligible to ∼1.6-fold more effective VN against R685S *vs. wt* at 7h. titers diminished for both viruses 20 h.p.i, but the effect was 4.9-fold higher for *wt* rVSV (**Supplementary Fig. 9b**).

Using A549 ACE2plusC3 cells, SARS-CoV-2 *wt* but not R685S generated syncytia between 7 and 20 h.p.i (**Supplementary Fig. 9c**). As previously reported, SARS-CoV-2 is more sensitive than rVSV-S to Ab mediated VN, likely related to the lower density of S on virions^44^. As with rVSV-S, *wt* rSARS-CoV-2 was slightly more resistant to VN at 7 h.p.i. potency dropped 2.5-fold on average for *wt* virus at 20 h.p.i, remaining constant for R685S virus (**Supplementary Fig. 9d**).

The resistance of *wt vs.* R685S to Ab-mediated VN, combined with the time-dependent decrease in Ab potency correlating with syncytia formation, supports the conclusion that syncytia formation enhances S escape from VN activity *in vitro* potentially contributing to FCS positive selection in immune hosts.

### S variants confirm syncytia-mediated IFN-escape

The rapid evolution of SARS-CoV-2 S in humans provides a golden opportunity to test the correlation between S-mediated syncytia formation and IFN-escape. The Delta S variant and its hallmark P681R substitution induce more and larger syncytia than ancestral S^3^. While the Omicron S variant is reported to reduce syncytia formation^45^, the prevalent P681H substitution in Omicron and Alpha S variants can increase syncytia^46^. To better establish the correlation between fusogenicity and IFN resistance, we generated S proteins from the Delta and Omicron variants and *wt* S variants with hallmark proximal FCS substitutions P681R and P681H.

We first assessed S mediated-syncytia formation using cells expressing S from a transgene-encoded mRNA linking S and GFP expression via an IRES. While there were comparable numbers of GFP-expressing transfectants, GFP-positive syncytia were significantly reduced in R685S and Omicron S variants compared to *wt* S (**Fig.2c** and **Supplementary Fig. 10a**). As expected, Omicron’s compromised syncytia generation was not simply attributable to the P681H substitution, which enhanced syncytia when introduced into *wt* S. Conversely, substituting P681R from Delta variant into *wt* S and Delta S itself increased syncytia 1.5- and 4-fold over *wt* S.

We extended these findings to rVSV-S infection of Vero cells pretreated with IFN-β (0.1 ng/ml) for 20 h. VSV expressing Delta or P681R S formed larger syncytia than VSV-*wt* S, with the opposite observed for Omicron S (**Fig. 5a** and **5b**). We repeated this experiment using type I (α and β), II (γ), and III (λ2) IFNs over a wide dose range to determine the IFN IC_50_ for blocking infection as measured by total eGFP reporter fluorescence 20 h.p.i. (**Fig. 5c**). Among five rVSV-S viruses tested, R685S was most sensitive to all three types of IFNs, followed by Omicron, *wt*, P681R and Delta, with this rank order maintained for each IFN tested. We calculated the fusion index (**Supplementary Fig. 10b**, number of nuclei present in syncytia/number of individual infected cells) for each IFN and plotted the fusion index against the IC_50_ for the five rVSV-S viruses tested (**Fig. 5d**). This reveals a highly robust correlation between syncytia formation and IFN IC_50_, with R^2^ values ranging from 0.8127 to 0.9543, consistent with causality.

**Fig. 5.**
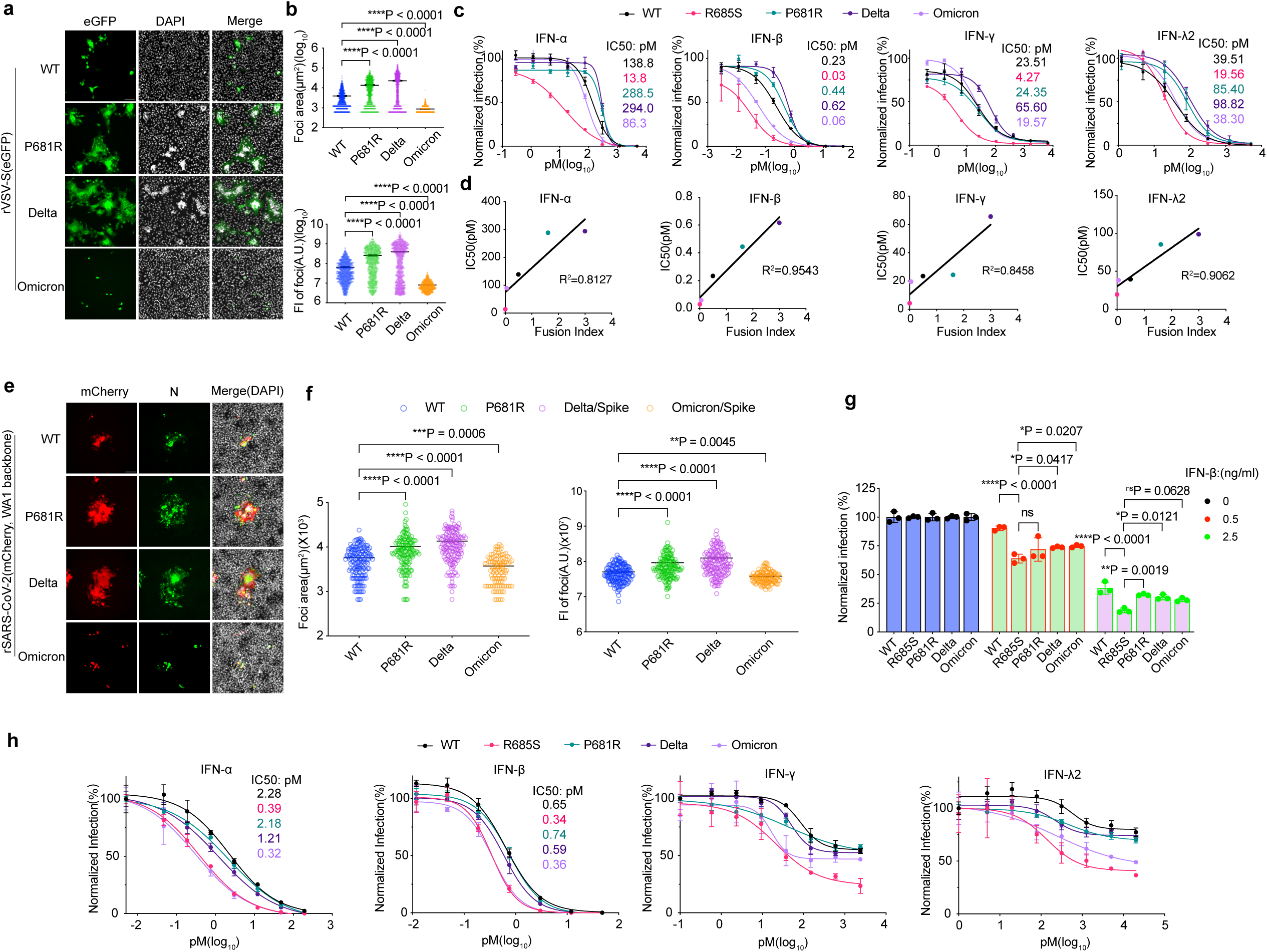
Correlation between syncytia formation and IFN resistance. **a** IFN-β (0.1 ng/ml) pretreated Vero cells were infected (MOI 0.01) with rVSV-S expressing P681R, Delta or Omicron (BA.1) S at 20 hrs post treatment. IFN-β (0.1 ng/ml) was maintained through infection. **b** As in **a**, with area (top) and fluorescence intensity (bottom) of individual infection foci measured following infection: *wt* (n=1147 foci), P681R (n=888 foci), Delta (n=1179 foci), Omicron (n=1004 foci). Data show mean ± s.e.m **c** Dose titration of indicated IFN-treated cells infected with rVSV- S virus expressing *wt*, R685S, P681R, Delta or Omicron S was determined in Vero cells at 20 h.p.i. Data were fitted using nonlinear regression. Data show mean ± s.d. (n=3 samples per dose). **d** Correlation of IC_50_ with fusion index for rVSV-S infected Vero cells. The fusion index was calculated by dividing the nuclei number of syncytia foci by the nuclei number of individual infected cells. Linear regression was performed to correlate the IC_50_ with the fusion index. Details are described in the Method section. **e** and **f** Vero cell infection 36 h.p.i with the rSARS-CoV-2 expressing S indicated. The area(left) and fluorescence intensity(right) of infection foci in the infection of rSARS-CoV-2 *wt*(n=156 foci), P681R(n=146 foci), Delta(n=149 foci), and Omicron(n=113 foci). Data show mean ± s.e.m. **g** IFN-β sensitivity of rSARS-CoV-2 virus bearing *wt*, R685S, P681R, Delta and Omicron S in Calu-3 cells at 24 h.p.i. Data show mean ± s.d (n=3 biological replicates). **h** Dose titration of rSARS-CoV-2 virus bearing *wt*, R685S, P681R, Delta and Omicron S against type I (IFN-α and −β), type II (IFN-γ) and type III (IFN-λ2) IFNs were determined in ACE2plusC3 cells at 24 h.p.i. Data show mean ± s.d (n=3 biological replicates). Statistical analysis was performed using a nonparametric Kruskal-Wallis test (**b,f**) or two-way ANOVA test(**g**) with multiple comparisons.

We repeated this experiment by infecting Vero cells for 36 h (MOI 0.05) with *wt* rSARS-CoV-2 virus and viruses with S replaced by one of the four S tested above. As with rVSV-S, in cells treated with IFN-β (0.1 ng/ml) relative to *wt* S P681R- and Delta-S viruses generated larger and more intense foci measured either by mCherry reporter expression or staining for SARS-CoV-2 nucleocapsid (N) while Omicron foci were smaller and less intense (**Fig. 5e** and **5f**). Using Calu-3 human lung epithelial cells treated with higher amounts of IFN-β (0.5 and 2.5 ng/ml), R685S was most sensitive to the IFN, followed by the Omicron. Interestingly, the *wt* showed the lowest sensitivity compared to the other four S variants (**Fig. 5g**). Lastly, in A549 ACE2plusC3 cells which has been shown to form strong syncytia upon SARS-CoV-2 infection^47^, *wt*-, P681R- or Delta-S rSARS-CoV-2 viruses exhibited robust syncytia formation, while R685S- or Omicron-S viruses displayed minimal syncytia (**Supplementary Fig. 10c**). R685S and Omicron viruses displayed similar higher sensitivity to IFN-β and IFN-αthan *wt*, P681R, and Delta (**Fig. 5h**). ACE2plusC3 cells were much less sensitive to IFN-γ or IFN-α2 against SARS-CoV-2 infection but showed a similar difference in S-dependent anti-viral activity (**Fig. 5h**). In summary, by examining five different S variants in two viral systems and three cell lines treated with four different IFNs, our results demonstrate a positive correlation between S-dependent syncytia and IFN escape.

### S fusogenicity of dominant circulating variants increases in each major evolutionary phase

How did SARS-CoV-2 evolution in humans over 3 years alter S fusogenicity? We examined S from nine dominant strains by mixing Vero cells transfected with a plasmid encoding complementing dual split proteins (DSP) DSP_1-7_ or DSP_8-11_(ref^48^) and transfected the mixed culture with S-expressing plasmids. We measured syncytia formation by the generation of fluorescent eGFP from DSP_1-7_ association with DSP_8-11_, quantitating the number of syncytia, syncytial area, and fluorescent intensity. This revealed increased S fusogenicity as WA1 evolved to D614G, Alpha, and Delta variants, with a decrease to sub-WA1 levels in the original Omicron strain (BA.1) with a steady fusogenicity increase with BA.2, BA4/5, BQ.1 and XBB variants (**Fig. 6a to c**).

**Fig. 6.**
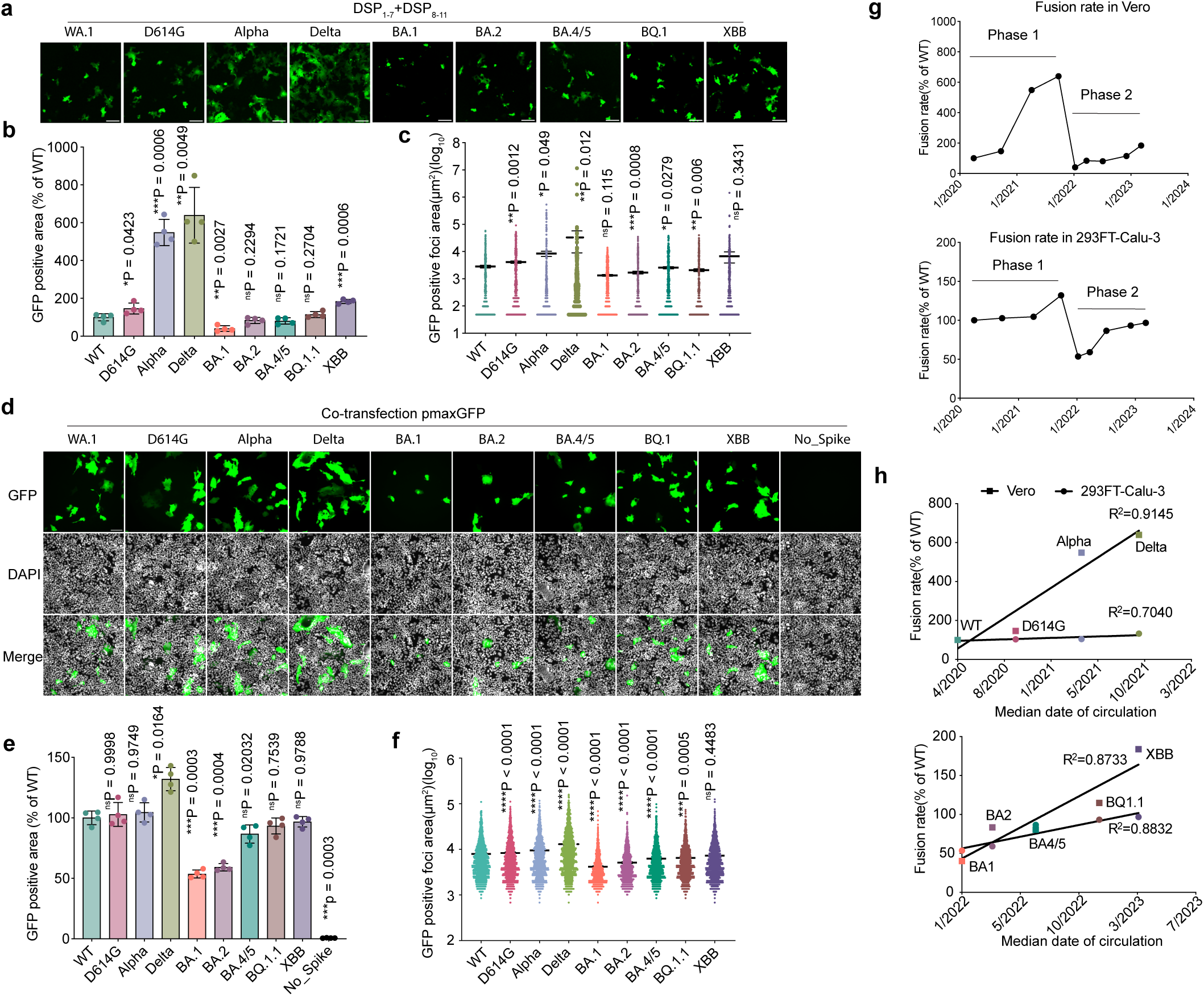
Fusogenicity over the course of SARS-CoV-2 evolution occurs in a phase-dependent manner. **a-c** Syncytia formation mediated by S protein from circulating SARS-CoV-2 variants was measured in Vero cells using the GFP dual split protein (DSP) system. Vero cells transfected with DSP_1-7_ were mixed with Vero cells transfected with DSP_8-11_, then the mixed cells were transfected with S-bearing plasmid. Representative images showing syncytia mediated by S from indicated circulating variants at 24 h.p.i. (**a**). Total GFP positive area normalized to *wt* S (**b**) and GFP positive individual focus area (**c**) were determined (n=4 wells per group). Ctrl Vec, Control Vector. DL, Detection Limit. **d-f** Syncytia mediated by S protein from circulating SARS-CoV-2 variants was measured in 293FT and Calu-3 cells. 293FT cells co-transfected with S-and pmaxGFP expression plasmids were overlayed on 100% confluent Calu-3 cells for 24 h. Total GFP positive area normalized to *wt* S (**e**) and GFP positive individual focus area (**f**) were determined (n=4 wells per group). **g** and **h** Correlation of fusion rate with median isolation date of each circulating variant in two phases. Two independent assays for measuring cell-cell fusion rate of each S of dominant circulating variants showed an increased fusogenicity of S in 2 phases (**g**). In **h**, the top shows the correlation of variants of *wt*, D614G, Alpha and Delta in phase 1, and the bottom shows Omicron subvariants in phase 2 (**h**). Statistical analysis was performed using a one-way ANOVA with multiple comparisons. *P < 0.05, **P < 0.01, ***P < 0.001 and ****P < 0.0001; ns, not significant.

We repeated this experiment using an alternative assay based on syncytia formation between human 293FT cells co-transfected with plasmids expressing pmaxGFP and S and human lung epithelial Calu-3 cells (**Fig. 6d**). This demonstrated the same pattern, though with smaller time-dependent increases in Alpha and Delta variant fusogenicity (**Fig. 6d to f**).

To rule out that apparent differences in fusogenicity are related to levels of cell surface expression of S, we measured surface (fixed cells) and total S protein expression (fixed and saponin permeabilized cells) of Vero and 293FT cells transfected with S-expressing plasmids using flow cytometry. To prevent syncytia formation in Vero cells—which endogenously express ACE2—we assessed surface and total S expression at 10 hours post-transfection before significant fusion could occur.

Interestingly, we found that while total S expression levels in Vero cells were comparable across variants, Omicron subvariants—particularly BA.1, BA.2, and XBB—exhibited higher surface expression of S protein (**Supplementary Fig. 11a**). We further assessed the kinetics of S expression in 293FT cells at 6, 10, and 24 hours post-transfection (**Supplementary Fig. 11b**). Consistent with the Vero cell data, Omicron subvariants displayed more rapid surface expression kinetics. Taken together with their low fusogenicity, these results suggest that surface S expression levels are not a major determinant of the spike protein’s fusogenic potential.

Interestingly, plotting fusogenicity *vs.* the median date of variant emergence (**Supplementary Fig. 12**) showed an increased fusogenicity of S following SARS-CoV-2 evolution in 2 phases (**Fig. 6g**). Moreover, linear regression analysis of S fusogenicity in each phase, with the median date, yielded a positive correlation for both assays, with R^2^ values ranging from 0.70 to 0.91 (**Fig. 6h**). These data suggest a new model for SARS-CoV-2 evolution, with newly introduced strains with major antigenic changes (WA and Omicron BA.1) with low fusogenicity evolving under adaptive and innate immune pressure to increase fusogenicity.

### P14-mediated syncytia formation promotes viral resistance to interferons and neutralizing antibodies

Extending our findings beyond S-mediated syncytia, we examined the role of syncytia formation induced by the fusion-associated small (125 residues) transmembrane (p14 FAST) protein from reptilian orthoreovirus—a non-enveloped viral protein—in antiviral responses. We assessed the effect of p14-mediated syncytia formation on viral replication by infecting transiently p14-transfected cells prior to widespread syncytial formation.

In p14-transfected Vero cells, we observed a modest but significant (∼1.5-fold) decrease in viral titers following VSV infection at 24 and 36 h.p.i. (**Fig. 7a**), consistent with results seen with SARS-CoV-2 S-mediated syncytia. We next evaluated the impact of p14-mediated syncytia on IFN resistance in Vero cells using reporter viruses VSV, influenza A virus (PR8), and seasonal coronavirus OC43 (**Fig. 7b**). After type I IFN treatment p14 expression enhanced VSV viral replication, as evidenced by larger infection foci (**Fig. 7c**), increased fluorescence intensity (2–3 fold), and elevated viral titers in the supernatant (3–7 fold) (**Fig. 7d and Supplementary Fig. 13a**). p14-enhanced interferon resistance conferred was also observed for PR8 and OC43 viruses (**Fig. 7d, f, and Supplementary Fig. 13b–e**). In all cases, infection of p14-induced syncytia was obvious from light microscopy.

**Fig. 7.**
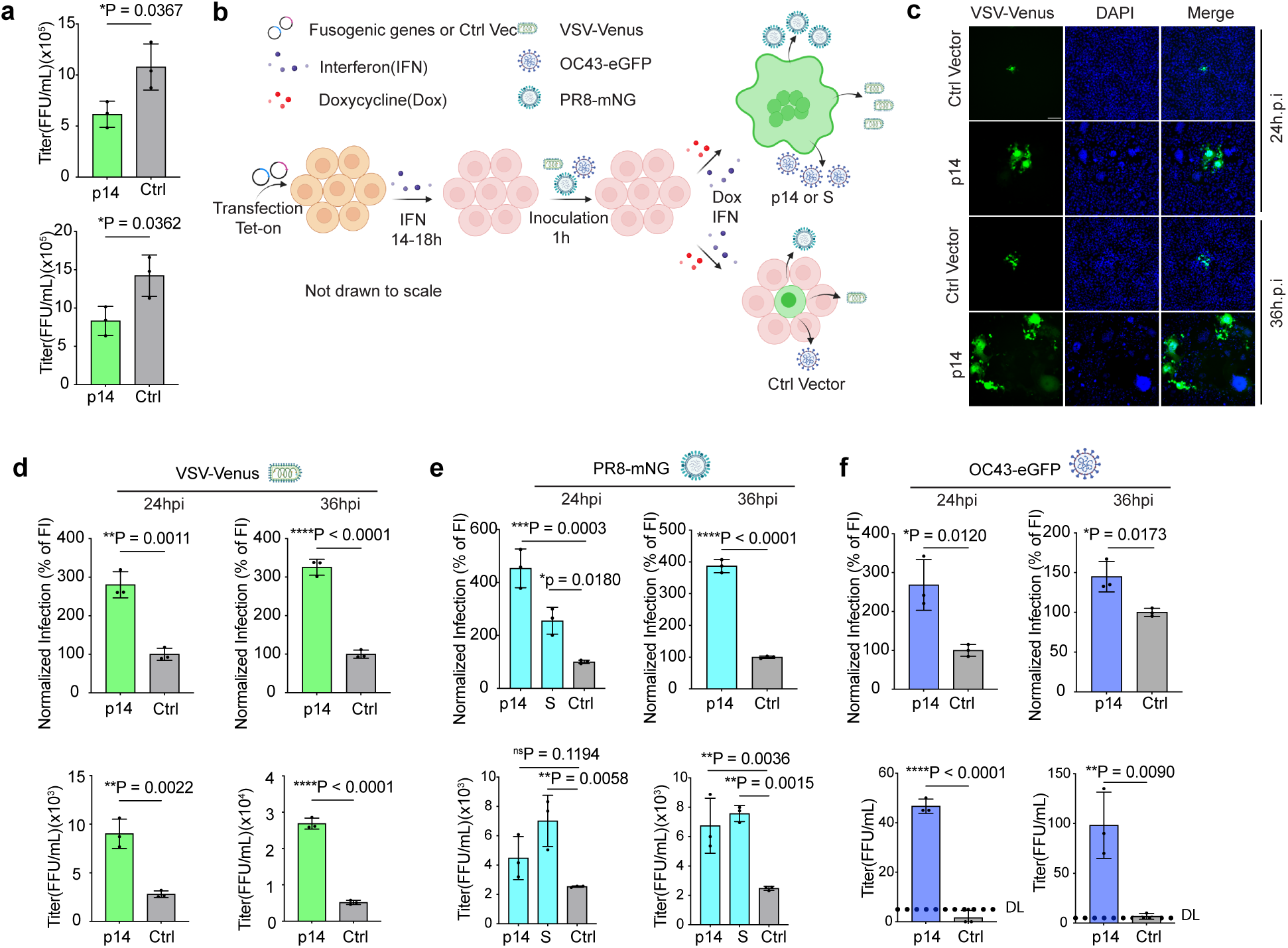
p14-mediated syncytia formation enhances virus resistance to IFN. **a** Effect of p14-mediated syncytia formation on viral production. Vero cells were transfected with a p14-expressing plasmid and infected with VSV at MOI of 0.1. Supernatants were collected at 24 h.p.i.(top) and 36 h.p.i. (bottom). Cell debris was removed by centrifugation, and viral titers were determined on Vero cells as described in the Methods section. n = 3 biological replicates. **b** Schematic diagram (Created in BioRender. Gibbs, J. (2026) https://BioRender.com/wpfype0) of the experimental design to assess the effect of p14-mediated syncytia formation on viral resistance to IFN in Vero cells. Ctrl Vec, Control Vector **c** Representative images of VSV infection in p14-mediated syncytia formation in Vero cell treated with 0.5ng/mL IFN-β. Scale bar, 100μm **d-f** Fluorescence intensity (top) and viral titers (bottom) in Vero cells with syncytia formation mediated by p14 or SARS-CoV-2 S protein under IFN-β treatment. Vero cells were pretreated with IFN-β and infected with recombinant VSV (**d**), PR8 influenza virus (**e**), or OC43 coronavirus (**f**) at MOI of 0.1. IFN-β concentrations of 0.5 ng/mL (rVSV), 1 ng/mL (rPR8), and 5 ng/mL (rOC43) were used. n = 3 biological replicates. DL, Detection Limit. Statistical analyses were performed using an unpaired t test (**a**, **d**, **e** (upper right panel), f) or one-way ANOVA with multiple comparisons (**e**, upper right panel excluded)). *P < 0.05, **P < 0.01, ***P < 0.001 and ****P < 0.0001; ns, not significant. Data show mean ± s.d.

We further investigated whether p14-mediated syncytia affect antibody neutralization. In VSV-infected cells, neutralizing monoclonal antibodies 1E9F9 and 8G5F11 inhibited over 95% of infection in non-syncytial conditions but only achieved 2–3 fold reduced efficacy in p14-mediated syncytia (**Supplementary Fig. 14a–c**). Similarly, a panel of monoclonal antibodies targeting Sa, Sb, Ca and Cb antigenic sites on PR8 effectively inhibited infection by 70–50% in Vero cells, but this neutralization was diminished by 2–3 fold in the presence of p14-induced syncytia (**Supplementary Fig. 14d, e**).

Together, these findings demonstrate that interferon resistance and antibody escape conferred by syncytia formation extend to many other viruses and may be a general factor in the evolution of fusogenic viruses.

## DISCUSSION

We examined why the FCS has been nearly perfectly maintained in millions of isolates during the first 3 years of SARS-CoV-2 evolution in humans. Previous studies established the critical role of the FCS in SARS-CoV-2 replication in human nasal epithelial cells and ferret transmission ^21^ and that an intact FCS increases pathogenesis in mice and hamsters ^20,22–25^.

Our findings support the conclusion that the FCS increases SARS-CoV-2 fitness by enhancing syncytia formation, enabling virus escape from virion entry inhibition mediated by IFN-induced proteins. In IFN-incompetent cells, *e*.*g.,* Vero cells which cannot secrete IFN, syncytia retard viral replication, resulting in the rapid selection of FCS loss mutants, as reported in many studies^20–22,49^. We show that syncytia enable S-expressing VSV to escape IFN, suggesting a general explanation for selecting syncytia-forming viruses in the presence of IFN pressure. We further demonstrate that syncytia formation reduces the efficiency of VN Abs *in vitro*, consistent with a contribution to FCS fitness in immune hosts. This may be a mass action effect based on the need for Ab to block more S present on infected cells *vs.* virions. Additionally, we show that following syncytia formation, recipient cell ribosomes are recruited to synthesize new viral proteins, likely increasing viral replication to partially mitigate the inherent disadvantage of cell-to-cell *vs.* virion-based virus infection.

SARS-CoV-2 S evolution in humans exhibits a punctuated pattern. Following the December 2019 SARS-CoV-2 introduction into humans, S evolved relatively gradually (though faster than IAV HA, the previous poster virus for antigenic drift). The appearance of Omicron BA.1 in Nov 2021 marked a major acceleration in SARS-CoV-2 evolution with 34 S amino acid substitutions compared to circulating strains, more akin to antigenic shift in IAV than antigenic drift. This virus likely evolved in an immunocompromised individual, enabling sequential selection of Ab escape mutants ^50–56^. If such an individual were also deficient in IFN-mediated anti-viral activity, this would greatly favor the selection of less fusogenic viruses, which we show have a selective growth advantage in IFN pathway-compromised cells.

While our study does not directly examine the association between fusogenicity and viral pathogenicity, the SARS-CoV-2 literature suggests a positive correlation. The hallmark substitution P681R in the Delta variant that augments fusogenicity contributes to pathogenicity^3^. The pathogenicity of Omicron BA.1, while attenuated compared to Delta, increases in parallel with fusogenicity with variant evolution^45,57–60^. Importantly, as SARS-CoV-2 continues to evolve, several recent Omicron subvariants—including XBB.1.5, EG.5, BA.2.86, and their descendants—have exhibited comparable or even enhanced fusogenicity of the S protein, particularly in the context of infection^61–63^. These findings suggest ongoing positive selection for syncytia formation during SARS-CoV-2 evolution. Occasional syncytia were observed in K18-hACE2 transgenic mice, and clinical reports have documented syncytia in over 70% to more than 90% of examined human lung specimens^17,64,65^. These observations indicate that cell-cell transmission may represent a significant mechanism of viral spread within the host, although further investigation is warranted.

In conclusion, our study highlights the essential role of the S FCS in promoting viral syncytia, allowing SARS-CoV-2 to evade innate and likely adaptive immunity. Notably, the IFN and antibody resistance conferred by syncytia formation was also observed in recombinant VSV, IAV (PR8), and OC43 viruses engineered to express SARS-CoV-2 S or the p14 FAST protein from a non-enveloped virus. This supports a broader model in which syncytia formation serves as a conserved mechanism for immune evasion across diverse fusogenic viruses. These findings underscore the importance of monitoring FCS acquisition and fusogenicity in viral evolution, particularly in the context of future epidemics or pandemics.

## METHODS

### Cell culture

We used baby hamster kidney fibroblast BHK21 (ATCC; CCL-10), African green monkey kidney Vero E6 (ATCC; C1008, Clone E6) Vero (provided by Dr. Nihal Altan-Bonnet, NHLBI), human embryonic kidney HEK293FT cells (Thermo Fisher; R70007), human lung adenocarcinoma A549 cells (ATCC; CCL-185) Calu-3 cells (ATCC; HTB-55), human colorectal adenocarcinoma Caco-2 cells (ATCC; HTB-37) cells. All cells were cultured in Dulbecco’s modified Eagle’s medium (DMEM) (Gibco; 10569044) supplemented with 10% fetal bovine serum (FBS, HyClone, no. SH30071.03) and 50 μg/ml Gentamicin (Quality Biological). We generated BHK21 cells stably expressing human ACE2 (BHK21-ACE2), A549 cells stably expressing human ACE2 (A549-ACE2), Vero E6 cells stably expressing human TMPRSS2 (Vero E6-TMPRSS2) and 293FT cells stably expressing T7 polymerase and VSVG (293FT-VSVG-T7pol) using the Sleeping Beauty transposon plasmid expression system ^66^. Vero E6 cells stably expressing human ACE2 and TMPRSS2 (Vero E6-AT2) were obtained from BEI (NR-54970). A549 cells stably expressing human ACE2 and TMPRSS2 (ACE2plusC3, ATCC; CRL-3560) were provided by Dr. Ching-Wen Chang (University of Massachusetts Chan Medical School). Cell lines were confirmed to be mycoplasma-free using MycoStrip (InvivoGen, rep-mys-50). Primary human small airway epithelial cells (HSAEC, ATCC, PCS-301-010) were purchased from ATCC and maintained in PneumaCult™-Ex Plus Medium (STEMCELL Technologies) according to the manufacturer’s instructions. All cell lines were incubated at 37 °C and 5% CO_2_ in a humidified incubator.

### Plasmid construction

The plasmid-based VSV reverse genetic system (pVSV eGFP dG) was obtained from Addgene (#31842) and modified to insert the wild-type *S* gene of the original Wuhan-Hu-1 strain of SARS-CoV-2 (GenBank MN908947.3) with a 21 amino acid deletion in its C terminal, between M and L genes, using standard molecular techniques. In brief, the VSV antigenome without the G gene and the SARS-CoV-2 *S* gene were amplified by PCR using Platinum™ SuperFi II PCR Master Mix (Thermo Fisher). PCR products were gel-purified and assembled using NEBuilder® HiFi DNA Assembly Master Mix (NEB). The assembled product was transformed into NEB 5-alpha Competent E. coli (NEB) using the standard protocol, and the cells were plated on carbenicillin LB agar plates (Quality Biological). Colonies were selected for Sanger sequencing to confirm the *S* gene insertion. Confirmed clones were cultured, and plasmids were isolated using the Midi prep plus kit (Qiagen). pVSV-SARS-CoV-2-S with mCherry was constructed using the same method by replacing eGFP in the antigenome.

We generated rSARS-CoV-2 viruses using the bacterial artificial chromosome (BAC)-based SARS-CoV-2 reverse genetic system^67^. The pBAC-SARS-CoV-2-mCherry-2A was digested with BstBI (FastDigest, Thermo Fisher) and BamHI (FastDigest, Thermo Fisher) and the larger fragment was purified by Zymoclean Large Fragment DNA Recovery Kit (Zymo Research). An intermediate plasmid, pUC57-orf1-S, containing a portion of the pBAC-SARS-CoV-2 genome, was used to introduce mutations in the FCS of the *S* gene or to generate Delta- and Omicron-S variants. To generate pUC57-orf1-S R685S, P681R, site-directed mutagenesis was performed on the pUC57-orf1-S plasmid. To create pUC57-orf1-S Delta-S and Omicron-S, the WA1 *S* gene was replaced by the Delta or Omicron *S* gene using NEBuilder® HiFi DNA Assembly Master Mix following the manufacturer’s instructions. Expected nucleotide sequences of pUC57-orf1-S with R685S, P681R, Delta or Omicron S gene were confirmed by Sanger sequencing. PCR reactions were conducted to amplify the fragment with partial orf1 and *S* genes, and products were assembled with the larger fragment by NEBuilder® HiFi DNA Assembly Master Mix. The assembled reaction was transformed into NEB® 10-beta Competent E. coli cells (NEB) following the manufacturer’s instruction. Bacterial colonies were selected for Sanger sequencing to confirm the *S* gene sequence, and confirmed colonies were subjected to maxiprep to isolate the BCA plasmids.

The sleeping beauty system^66^ was obtained from Addgene. Briefly, the genes for hACE2, TMPRSS2, codon-optimized T7 polymerase or VSVG were PCR amplified using primers containing BstBI restriction enzyme site. PCR products were purified and subcloned into pSBbi or tet-on inducible pSBtet vector that had been digested by BstBI. Confirmatory sequencing was performed on all constructs.

To express S we used a pHAGE-eGFP backbone vector. Briefly, genes encoding S and S variants were PCR amplified and subcloned into pHAGE-eGFP, forming an expression cassette of eGFP-IRES-S using NEBuilder® HiFi DNA Assembly Master Mix. All *S* genes lacked codons for the 21 C-terminal residues to increase cell surface expression. Codon-optimized full-length S-bearing pCAGGS plasmids, except the one containing *wt* S, were obtained as a gift from Marceline Côté at Addgene. wt S was generated by standard site mutagenesis from D614G S. All plasmids were sequenced to confirm nucleotide sequence fidelity.

### Generation of stable cell lines

Stable cell lines, except as indicated, were generated by the Sleeping Beauty transposase system. Briefly, cells were seeded on a 60-mm plate, and the following day, cells were co-transfected with 0.5 μg of pCMV(CAT)T7-SB100 (transposase vector; Addgene, 34879) and 5 μg of pSBbi or pSBtet containing gene of interest using TransIT-LT1 transfection reagent (Mirus Bio), following the manufacturer’s instructions. After 24 h, cells were detached and transferred to T-75 flasks, followed by selection with corresponding antibiotics for 2 weeks. Cell surface expression of hACE2 or TMPRSS2 was confirmed by flow cytometry. The functionality of T7 polymerase and VSVG expression in 293FT-T7pol-VSVG cell lines was determined using pUC19-T7pro-IRES-EGFP (Addgene, 138586) or through the recovery of rVSV virus.

### Generation of rVSV-SARS-CoV-2 S

Due to the lower efficiency of the rVSV system for rescuing viruses^68^, we optimized the rescue system using a stable 293FT cell line expressing T7 polymerase and VSVG, without using vaccina helper virus vTF7-3. This optimized system allowed us to efficiently recover rVSV-S, bearing an ancestral SARS-CoV-2 *wt* S as well as S variants. The generation of rVSV-S and its use in tissue culture at biosafety level 2 conditions was approved by the Institutional Biosafety Committee (IBC) at The National Institutes of Health (NIH)(IBC approved case number: RD-20-IV-23). Plasmid-based rescue of the rVSV was performed as described ^30,31^ with modifications. Briefly, 293FT-T7pol-VSVG cells in a 6-well plate were transfected with the VSV antigenome plasmid, along with plasmids expressing codon-optimized T7 polymerase (T7opt in pCAGGS, addgene, 65974), and VSV -N, -P, -L and -G-expressing plasmids (Addgene, 64087, 64088, 64085 and 8454), using TransIT-LT1 transfection reagent. Media were exchanged with fresh complete DMEM containing doxycycline (1μg/ml, for VSVG expression). Cells were monitored by fluorescence microscopy every day for 2-3 d. Transfected cells with GFP- or mCherry-positive clusters and typical cytopathic effect (CPE) were collected as passage 0 and inoculated into fresh BHK21-ACE2 cells to propagate viral stock P1, which underwent further propagations in BHK21-ACE2 cells. Viral passages were subjected to Sanger and next-generation sequencing to verify *S* gene or viral genome sequences. Plaque purification was performed on Vero cells with viral passage 4. Each purified plaque was Sanger sequenced for the entire *S* gene. Plaque-purified virus, along with rescued rVSV viruses bearing P681R, Delta, and Omicron S, were propagated in Vero E6-TMPRSS2 cells. Supernatants were confirmed by NGS, aliquoted, and stored at −80°C for further experiments.

### Generation of recombinant SARS-CoV-2

We fused the mCherry gene with the N gene via a 2A linker using a bacterial artificial chromosome-based SARS-CoV-2 reverse genetic system^67^. Confluent BHK21-ACE2 (2×10^6^ cells/well in 6-well plates, duplicates) were transfected with 2.5 μg/well of pBAC-SARS-CoV-2 using TransIT-LT1 transfection reagent. Media were exchanged with fresh DMEM containing 2% FBS 6 h post-transfection. After 48-72 h, mCherry-positive cells displaying typical viral infection were detached and collected, along with supernatant, labeled as P0, and stored at −80°C. The P0 viral stock was centrifuged to remove cell debris and used to infect fresh Vero E6-TMPRSS2 cells for 48-72 h. The supernatant was collected as P1. After confirming the rescued virus (P1) by Sanger sequencing, the P0 virus underwent two rounds of propagation, resulting in a new virus stock (P2), which was titrated for further experiments after NGS confirmation of the viral genome sequence. No additional unexpected mutations that could interfere with the IFN signaling pathway were identified (**Supplementary Table 1**).

### Virus infectivity titration

To titrate virus, Vero E6 or Vero E6-TMPRSS2 cells were seeded in a 12-well plate at 3×10^5^ cells per well. The following day, viral stocks were serially diluted and inoculated onto confluent cells for 1 h with gentle shaking every 15 min. After removal of the inoculation, the cells were covered by 1× plaque MEM medium containing 4% FBS and 1.25% Avicel. At 24 h.p.i, cells were fixed with 4% paraformaldehyde (PFA) or 10% neutral buffered formalin (NBF) for 30 min at RT. The fixed plates were scanned using the high-content imaging system Cytation 5 (Agilent), with either the GFP channel for rVSV-S or the Texas red channel for rSARS-CoV-2 virus. Raw images were processed and stitched with Gen5 3.12 (Agilent) in a default setting, and focus-forming unit (FFU) was determined.

For titration of viral stocks collected from infected monolayer cells or ALI apical wash, Vero E6-TMPRSS2(for monolayer) or Vero E6-AT2 (for ALI Apical wash) cells were seeded in a black 96-well plate at 3×10^4^ cells per well. The following day, viral supernatant was serially diluted and inoculated onto the cells for 1 h with gentle shaking every 15 min. After removal of the inoculation, fresh completed DMEM medium was added and incubated for 7-8 h. Following incubation, cells were fixed with 4% PFA or 10% NBF for 30 min at RT. The fixed plate was scanned by high-content imaging system Cytation 5 with either GFP channel for rVSV-S or Texas red channel for recombinant rSARS-CoV-2 virus. The raw images were processed and with Gen5 3.12 in a default setting, and FFU were determined using the cellular analysis mode. Alternatively, cells were detached by TrypLE™ Express Enzyme (Thermo Fisher), fixed by 4% PFA for 30 min at RT, and subjected to flow cytometry analysis.

### RNA extraction, RT–PCR, and Sanger sequencing

Viral RNAs were extracted by the QIAamp Viral RNA Mini Kit (Qiagen) according to the manufacturer’s instructions. After extraction, the RNAs were dissolved in 20 μl nuclease-free water. Two μl of RNA samples was used for reverse transcription with the AccuScript High-Fidelity 1st Strand cDNA Synthesis Kit (Agilent; 200820) using random hexamer primers. DNA fragments containing the entire *S* gene were amplified by PCR. The resulting DNAs were purified by the QIAquick PCR Purification Kit (Qiagen), and the sequences were determined by Sanger sequencing by Psomagen (Rockville, MD).

### Next-generation sequencing

Ten ml viral stock total RNA was used as input for NGS library preparing following the Illumina Stranded Total RNA Prep, Ligation (Illumina). Purified libraries were quantified using the Kapa Library Quantification Kit (Roche), pooled in equimolar concentrations, and sequenced as 2 x 150 bp reads on the MiSeq instrument using the MiSeq Reagent Micro kit v2 (Illumina). Raw image files were converted to fastq files using bcl2fastq (v2.20.0.422, Illumina) and processed as previously described ^69^. Reference sequences used for mapping included SARS-CoV-2 genome (MN985325.1), Delta and Omicron S sequences, as well as the rVSV-S construct sequence. Detected variants were confirmed by visual inspection using the Integrative Genomics Viewer^70^.

### Cultured Cell Infection

We infected Vero or ACE2plusC3 cells with rVSV-S or rSARS-CoV-2 as described ^20^. In brief, cells were seeded in a 24-well plate one day prior to infection. on the following day, confluent cells were infected with rVSV-S or rSARS-CoV-2 at a MOI of 0.01 (or indicated MOI) at 37 °C for 1 h with gentle shaking at every 15 min. Cells were washed twice with DPBS and fresh complete medium was added. Supernatant was collected and centrifugated to remove cell debris and stored at −80 °C. Infected monolayers were fixed with 4% PFA for 30 min at RT and scanned by a Cytation 5 and fluorescence intensity was determined by cellular analysis mode in Gen5 3.12. For time series measurement of replication kinetics (Supplementary Fig. 3a-c), 100 FFU of each virus was applied to Vero cells in a 12-well plate. Images were captured at 24, 28, 36 and 48 h.p.i. Area and fluorescence intensity of infection foci were determined in cellular analysis mode in Gen5 3.12.

### Western blotting

Vero cells infected with rVSV-S (wild-type or R685S mutant) were lysed using RIPA buffer (Thermo Scientific) supplemented with a protease inhibitor cocktail (cOmplete™, Roche). Equal volumes of each sample were loaded onto 4–12% Bis-Tris Mini Protein Gels (NuPAGE™, NP0322BOX) and separated by SDS-PAGE. Proteins were then transferred to nitrocellulose (NC) membranes using iBlot 2 Gel Transfer Device. The membranes were probed with SARS-CoV-2 S2 subunit-specific antibodies (R&D Systems, MAB10557), followed by incubation with an IRDye® 800CW goat anti-mouse IgG secondary antibody (LI-COR, 926-32210). Protein bands were visualized using the Odyssey® DLx Imaging System (LI-COR), and band intensities were quantified using Image Studio software (LI-COR).

### IFN-β inhibition of infection

Vero cells were pretreated with the indicated concentration of IFN-β (Peprotech; 300-02BC) for 18-20 h. Subsequently, cells were inoculated with either rVSV-S or rSARS-CoV-2 for 1 h. After removing the inoculum, fresh media with the indicated concentration of IFN-β was added. Infected cells were either scanned live or after PFA fixation (4%) at indicated times. The fluorescence intensity of infection foci was determined using cellular analysis mode by Gen5 3.12.

For time series recording of rVSV-S infection in Vero cells under IFN-β treatment following the procedure described above cells were inoculated with WT and R685S at MOI 0.01 or indicated MOI for 1 h. Following inoculation, cells were washed, and fresh media with the corresponding concentration of IFN-β were added. At 18 h.p.i, plates were time-lapse recorded for eGFP channel using a 10× objective with Cytation 5 under 5% CO_2_ at 37°C. The interval time was 30 min for the recording. The movie (supplementary Movie.1-4) was made by Gen5 3.12 by default setting.

### Ruxolitinib treatment

A549-ACE2 cells were seeded in 24-well plate one day prior to infection. On the following day, the cells were treated with 2 μM ruxolitinib for 2 h and washed with PBS. Cells were then infected with *wt* or R685S virus with MOI of 0.05 for 1 h, washed, and fresh media with 2 μM ruxolitinib were added. Cells were then incubated for 48 or 72 h. The supernatant was centrifuged to remove cell debris and stored at −80°C. Infected monolayers were fixed by 4% PFA and scanned using a Cytation 5. Raw images were acquired, processed, and stitched with Gen5 3.12 by default settings. Cellular analysis mode was conducted to analyze the fluorescence intensity.

### Competition assay

Competition assays were performed as described ^20^. Ratios (50:50, 90:10, and 10:90 of *wt*: R685S) were determined by FFU derived from viral stocks. Cells were infected at an MOI of 0.1 (*wt* and R685S) as described in “Cultured Cell Infection” section. Infectivity titers were determined by flow cytometry as described above.

### IFN-β viral infection inhibition

Vero cells were seeded in a black 96-well plate at a density of 3 × 10^4^ cells per well and incubated overnight. Cells were treated with IFN-β at 4-fold serial dilutions for 20 h and infected with *wt* or R685S at an MOI of 0.05 for 1 h. Following inoculation, cells were washed, and fresh media were added to cells for incubation for 7 h at 37°C. Cells were then fixed with 4% PFA, and images were acquired using Cytation 5. Cellular analysis was performed to quantitate infections using Gen5 3.12. “Infection inhibition” was calculated by dividing the fluorescence intensity of IFN-β-treated samples by mock-treated samples.

### Transfection-based cell-cell fusion assay

Vero cells were seeded in a 24-well plate at a density of 1.5 × 10^5^ cells per well. The following day, receptor cells were transfected with pLifeAct-mScarlet. Donor cells were transfected with pHAGE-eGFP-IRES-S *wt* or R685S for 4-6 h and detached using trypLE. Detached donor cells were overlaid onto receptor cells at a 1:4 ratio for 20 h. Subsequently, cells were washed and fixed with 4% PFA. Images were acquired, and fluorescence intensity was determined as above.

### Emetine syncytia assay

We treated recipient Vero cells in a 12-well plate with IFN-β at the indicated concentrations for 20 hours. We infected donor Vero cells with rVSV-S wt or R685S for 8 hours at an MOI of 0.5. Then treated cells with 10 μM emetine (an irreversible protein synthesis inhibitor) diluted in complete medium at 37°C for 30 minutes. We detached cells with enzyme-free cell dissociation buffer (ThermoFisher, 13151014) and washed cells extensively in complete medium with medium changes every 15 minutes for 90 minutes to completely remove unbound emetine, which otherwise would be transferred to recipient cells. We overlaid donor cells on recipient cells at a 1:5 ratio, adding fresh medium containing the corresponding concentration of IFN-β. Cells were centrifuged for 5 minutes at 300 × g, and were carefully moved to an incubator. After 24 h, we carefully washed cells to remove floating donor cells prior to fixing with 4% PFA. Raw images were acquired using Cytation 5 and processed and analyzed with Gen5 3.12 software.

### Fluorescent virus microneutralization assay

VN assays were conducted using rVSV-S or rSARS-CoV-2 virus, following previously established protocols^20^. Vero-ACE2 cells were plated on a black flat-bottom 96-well plate (Costar, 3603). The next day, mAbs were serially diluted in 4-fold dilutions and incubated with 500 FFU of rVSV-S *wt* or R685S expressing eGFP at 37°C for 30 min and the virus-antibody mixture was transferred to cells. After 7 h, the plate was sealed with a parafilm membrane and scanned for eGFP fluorescence on Cytation 5 under 5% CO_2_ at 37°C. Subsequently, the plate was returned to the incubator at 5% CO_2_ and 37°C. At 20 h.p.i, the plate was re-scanned for eGFP fluorescence using the same settings as at 7 h.p.i.

A similar experimental procedure was conducted for the neutralization assay for rSARS-CoV-2. Due to BSL-3 laboratory limitation, a duplicate plate of each infection was set in the rSARS-CoV-2 infection. Plates were fixed with 4% PFA for 30 min at 7 and 20 h.p.i, respectively.

Raw images (2 × 2 montage) for each well were acquired using a 4× objective, processed, and stitched using default settings. eGFP-positive or mCherry-positive objects were quantified for each well. VN activity was determined by dividing the eGFP or mCherry intensity of mAb treated cells by the intensity of correspond mock-treated cells. The nonlinear regression fit (dose-response model) was employed to determine the neutralization dose 50% of eGFP or mCherry fluorescence (ND_50_).

### IFN inhibition titration

Vero or ACE2plusC3 cells were seeded in a black flat-bottom 96-well plate (Costar, 3603). The following day, cells were treated with varying doses of recombinant human IFN-α2 (Sino Biological; 13833-HNAY), IFN-β (Peprotech; 300-02BC), IFN-γ (Peprotech; 300-02), or IFN-λ2 (Peprotech; 300-02K) for 18-20 h before infection. Media were replaced with viral infection medium the next day, and fresh media containing IFN was added after 1 h inoculation. Cells were fixed with 4% PFA at RT for 30 min 20- or 24-h post-infection. Raw images of infected monolayers were acquired using Cytation 5. IFN inhibitions was calculated by ratio of fluorescence intensity of infection with IFN *vs.* mock treatment.

### Fusion Index calculation

For calculating the fusion index in rVSV-S infection in Vero cells, infected and non-infected (mock infected) cells were permeabilized and counterstained with DAPI for 10 min at RT. Raw montages (3×8=24) of infected cells were captured in DAPI and GFP channels using the high-content imaging system Cytation 5 with a 20x objective. The aggregation of nuclei (as shown in Fig. 4a,e) in syncytia making it difficult to count the nuclei number in syncytia directly. To do that, we first counted DAPI stained nuclei in mock infected cells using Gen5 to determine the total number of cells which represents the total number of cells in infection (N _cell total_). We determined the number of non-infected cells (N _cell no-infection_) in infected wells by using a GFP background level determined from non-infected cultures. We determined the number of non-syncytial cells (N _cell single_), including single infected and non-infected cells, based on the area of the DAPI signal (Gen5 setting: object size < 400 μm², the area of a single nucleus ranges from about 250-360 μm² *in situ*). The fusion index was then calculated using the following formula:

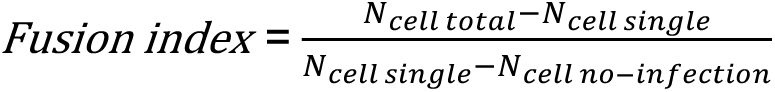

### Human respiratory epithelial cells in air-liquid interface format and infection

Human small airway epithelial cells were obtained from ATCC (PCS-301-010™, 3 donors: Lot Number: 70034740, 70035986, 70036650). After two passages in cell proliferation media (PneumaCult-Ex Plus medium, STEMCELL Technologies) supplemented with antibiotics, cells were seeded on Transwell inserts (0.4-micron pore size, 6.5mm, Corning; 3470) at a density of 33,000 cells per insert with media added to both the basal and apical sides. Once cells reached confluence, media were replaced with PneumaCult-ALI-S medium (STEMCELL Technologies) in the basal chamber, and the apical surface was exposed to establish an air-liquid interface (ALI). Monolayers were cultured at ALI for 4 weeks to promote differentiation into small airway epithelium. Nasal epithelium ALI was purchased from Epithelix (Plan-les-Ouates, Switzerland), which was differentiated from pooled cells of 14 donors (Batch Number: MP0011, differentiated on February 17^th^,2023; Experiments were performed in June 2023.), and was maintained in MucilAir™ medium (Epithelix).

For SARS-CoV-2 infection, differentiated ALI cultures were pretreated on the basal side with human IFN-β or IFN-λ2 at 0.2 or 10 ng/ml, respectively, for 20-24 h. Control wells were mock-treated with PBS. Following treatment, the apical side of the ALI was washed twice with DPBS for 5 min at 37°C to remove mucus. Cells were then inoculated with rSARS-CoV-2 *wt* or R685S virus at 10^4^ FFU diluted in 100 μl of plain DMEM at the apical side for 2 h at 37°C. After inoculation, the apical side was washed twice with DPBS and exposed to an air-liquid interface. The culture medium in the basal chamber was replaced with fresh IFN at 2 d.p.i, and 15 μl of fresh medium containing the corresponding IFN were added to the apical chamber to prevent drying after mucus removal, which would reduce cilia beating frequency. At 4 d.p.i., apical washes with DPBS for 20 min at 37°C were collected for further titration. Infected cells in the transwell were fixed with 4% PFA for 60 min before removal from the BSL-3 containment area.

### Immunofluorescence staining

The following primary Abs and IF dilutions were used in this study: Rabbit polyclonal anti-SARS-CoV-2-N (GeneTex, GTX135357, 1:500), rabbit monoclonal Alexa Fluor® 647 conjugated anti-Cytokeratin 5 (Cytk5, Abcam, ab193895, 1:200), rabbit monoclonal anti-Acetyl-α-tubulin (AcTub, Cell Signaling, 5335, 1:1000), mouse monoclonal Ab anti-Mucin 5AC (Muc5A, Invitrogen, MA5-12178, 1:100). The following secondary Abs were used in this study: cross-absorbed Goat anti-rabbit Alexa Fluor 750 or 488 (Invitrogen, A21039/A11008, 1:1000), cross-absorbed Goat anti-mouse Alexa Fluor 488 (Invitrogen, A11029, 1:1000).

For antigen detection, rSARS-CoV-2 infected epithelia were washed in DPBS and permeabilized with 0.5% TritonX-100 in DPBS for 20 min at RT and blocked with IF buffer (DPBS containing 5% normal Goat serum and 0.1% TritonX-100) for 1 h at RT after extensive washing with DPBS. Primary Abs were diluted in antibody dilution buffer (DPBS containing 1% BSA and 0.1% TritonX-100) and applied to inserts for 1 h at RT or overnight at 4°C. After three washes with DPBS, samples were incubated with secondary Abs for 1 h at RT. In the case of the primary antibody (Anti-Cytk5-AF647) derived from the same species as anti-AcTub, sequential staining was performed. Samples were incubated with 2 μg/ml of DAPI for 10 min at RT. Inserts were then washed extensively in DPBS, and membranes were removed from inserts and mounted in VECTASHIELD® Antifade Mounting Media (Vector Laboratories) prior to microscopy analysis.

### Microscopy and image analysis

For full ALI epithelial overviews, high-content imaging was performed using the Cytation 5 system with a 4X objective in a 24-well black glass bottom plate (Cellvis). Raw images were processed and stitched by default settings. Cellular analysis mode with object size ranging from 5 to 1000 μm was performed to quantify the overall fluorescence intensity of infection by Gen 5 3.12.

Ab stained infection foci were imaged using a Leica Stellaris 8 (Leica Microsystems, no. 11513859) in Z-stack mode, and three-dimensional (3D) reconstructions were done with Imaris in 3D view mode. The entire epithelia were tile scanned using a 63X 1.4 oil objective in low resolution with the LAS-X Navigator function. This entailed scanning 40×40 fields at 512×512 pixel resolution (1600 fields in total) for the mCherry channel. Subsequently, 20-45 random fields of infection were selected for Z-stack scanning, covering five channels (DAPI for nuclei, Alexa Fluor 488 for Muc5A, mCherry for rSARS-CoV-2 infection, Alexa Fluor 647 for Cytk5, and Alexa Fluor 750 for AcTub) for each epithelium. Images were captured at 1024 x 1024 pixel resolution with a step size of 0.25 μm. Representative images were processed by LAS-X software or Imaris (Bitplane).

To quantify syncytia (nuclei), randomly acquired fields were reconstructed into 3D structures, and individual planes were analyzed with Leica LAS-X software. The mCherry signal was used as a marker of infected cells, and the number of nuclei within each infected focus was determined by eye.

### Mouse infection

B6.Cg-Tg(K18-ACE2)2Prlmn/J (K18) transgenic mice (JAX Stock No. 034860) were bred at FDA White Oak Vivarium or purchased from the Jackson Laboratory. Homozygous K18 mice of both sexes at approximately 8−12 weeks were inoculated with rSARS-CoV-2 *wt* or R685S at 2000 FFU/50 μl/mouse under light isoflurane anesthesia. One day before and one day post infection, mice were treated i.n. with recombinant murine IFN-λ2 (Peprotech cat no. 250-33) at 2 μg/50 μl/mouse or 50 μl/mouse of PBS (mock-treated). Three d.p.i, mice were euthanized, and whole lungs were harvested for viral infectivity/PCR titers and cytokine determination. Procedures were performed according to the animal study protocols approved by the FDA White Oak Animal Program Animal Care and Use Committee. For histological and pathological analysis, including the detection of syncytia formation, female K18-hACE2 transgenic mice were intranasally inoculated with 10,000 FFU of recombinant SARS-CoV-2 *wt* or R685S mutant virus in a 50 µL volume. At 4 h.p.i., mice received intranasal treatment with either recombinant murine IFN-λ2 or PBS as a control. Body weight was monitored daily, and mice were euthanized at the indicated time points. Whole lungs were collected for histological examination and immunohistochemistry. Left brain was collected for viral titration and right brain was collected for histological examination and immunohistochemistry.

### Determination of lung viral titers

Lung viral titers were measured using both real-time PCR ^71–73^ and focus-forming assay (FFA) ^74^. RNA was extracted from whole lung homogenates using RNeasy Plus Mini Kit (Qiagen #74136) and converted to cDNA using the High-Capacity cDNA Reverse Transcription Kit (Thermo Fisher Scientific #4368813). Copy numbers of SARS-CoV-2 nucleocapsid (N) gene in lung tissues were determined using 2019-nCoV RUO Kit (Integrated DNA Technologies #10006713 and QuantiNova SYBR Green PCR kit (Qiagen #208052) according to the following cycling program: 95°C for 120 s, 95°C for 5 s (50 cycles), and 60°C for 18 s ^71–73^. Results were calculated based on a standard curve constructed using threshold cycle (Ct) values of serially diluted pCC1-CoV2-F7 plasmid expressing SARS-CoV-2 N. A value of 1 was assigned if gene copies were below the detection limits. FFA was also conducted to measure infectious viral particles in lung homogenates^74^. Supernatants of lung homogenates were serially diluted in MEM with 2% FBS and 1% antibiotics and were added to Vero E6-AT2 pre-seeded in 96-well tissue culture plates. Following the incubation at 37 °C and 5% CO_2_ for 1 h, unattached viruses were removed, and cells were overlaid with 1.2% Avicel (DuPont) after 1:1 (v/v) mixing with 2x EMEM with 4% FBS and 2% antibiotics ^74^. Cells were incubated at 37 °C and 5% CO_2_ for up to 24 h. After fixation, viral foci were detected using Cytation 5.

### Multiplex proinflammatory cytokine measurement

Proinflammatory cytokines in mouse lung homogenates were measured using a V-PLEX Proinflammatory Panel 1 Mouse Kit (Meso Scale Discovery # K15048D) according to the manufacturer’s instructions. Data were acquired in a MESO QuickPlex SQ 120 imager equipped with MSD Discovery Workbench 4.0.12 (LSR_4_0_12).

#### Histology and immunohistochemistry

Histology and immunohistochemistry were performed as described previously^75^, with some modifications. Briefly, mice were euthanized and perfused transtracheally with10% NBF, tissues were fixed in 10% NBF for 3 days prior to removal from the BSL3 for processing. Fixed tissues were sent to VitroVivo Biotech (Rockville, MD) for routine process, embedding in paraffin and section at 4 μm thickness. Sequential sections were stained with hematoxylin and eosin or stained for SARS-CoV-2 nucleocapsid using a monoclonal anti-SARS-CoV nucleocapsid antibody. For Immunohistochemistry (IHC), slides were deparaffinized in xylene and rehydrated through a graded series of decreasing ethanol concentrations. Antigen retrieval was performed by microwave boiling in 0.1 M sodium citrate buffer (pH 6.0) for two cycles (100% power for 2.5 minutes, followed by 20% power for 15 minutes). Slides were then washed in TBST (2 × 5 minutes) with gentle agitation. Endogenous peroxidase activity was quenched by incubating the sections in 0.3% hydrogen peroxide (H₂O₂) in TBS at RT for 15 minutes. Permeabilization was carried out using 0.025% Triton X-100 for 5 minutes, followed by TBST washes (3 × 5 minutes) with gentle agitation. Non-specific binding was blocked with Background Punisher (BioCare Medical, Pacheco, CA, USA) for 10 minutes. Sections were then incubated overnight at 4°C with 2 µg/mL rabbit anti-SARS-CoV-2 nucleocapsid monoclonal antibody [HL344] (GTX635679, GeneTex), diluted in Renoir Red Diluent (BioCare Medical). After additional TBST washes (3 × 5 minutes), sections were incubated at RT for 30 minutes with HRP-labeled Polymer Anti-Rabbit antibody (8114s; Cell Signaling, Danvers, MA, USA). Color development was performed using DAB Peroxidase Substrate (SK-4100; Vector Laboratories, Burlingame, CA, USA) for 2 minutes (brain) or 5–8 minutes (lung) at RT. Slides were counterstained with hematoxylin and imaged using a ZEISS Axioscan 7 microscope (Carl Zeiss Inc., Oberkochen, Germany). Mouse pathology evaluation was performed as described previously^76^.

#### Spike variants mediated fusion assay

One T75 flask of Vero cells was transfected with a DSP_1-7_ expression plasmid while another T75 Vero cells was transfected with a DSP_8-11_ expression plasmid. Four h post-transfection cells were detached using TrypLE and mixed at 1:1 ratio and in 24-well plates. The following day, mixed cells were transfected with pCAGGS-SARS-CoV-2 S plasmids expressing codon-optimized full-length S or S variants. 24 h post-transfection, cells were washed and fixed by 4% PFA for 20 min at RT before being subjected to high-content imaging.

For cell-cell fusion in 293FT and Calu-3 cells, 293FT cells were co-transfected with pmaxGFP and pCAGGS-SARS-CoV-2 S plasmids. Control samples were co-transfected with pmaxGFP and pCAGGS empty plasmids. Four h post-transfection, 293FT cells were detached and overlayed onto 100% confluent Calu-3 cells at a ratio of 1: 50. After 24 h, cells were washed and fixed with 4% PFA for 20 min at RT before being subjected to high-content imaging. Raw images were processed and stitched by default settings. Cellular analysis mode was performed to determine the overall fluorescence intensity and area of GFP-positive foci.

#### p14-mediated syncytia formation and viral resistance to interferon

The fusion-associated small transmembrane (FAST) protein p14 from Reptilian orthoreovirus (Addgene plasmid #50025) was cloned into a Sleeping Beauty transposon system for expression. Vero cells were seeded in 12-well plates and transfected the following day with plasmids encoding either p14 or SARS-CoV-2 S protein using a standard transfection protocol. At 4–6 hours post-transfection, the medium was replaced with fresh culture medium containing either vehicle control or IFN, and cells were incubated overnight. The next day, cells were infected with either reporter Vesicular Stomatitis Virus (VSV), influenza A virus (PR8), or human coronavirus OC43 at MOI of 0.1 for 1 h. Following viral adsorption, the inoculum was removed and replaced with fresh medium containing the same IFN treatment conditions and doxycycline (Dox, 0.5 µg/mL) to induce transgene expression. Supernatants were collected at the indicated time points for viral titration, and cells were fixed with 10% NBF for imaging. High-content imaging was used to visualize syncytia formation and infection levels. Viral titers in the supernatants were determined as described in the “Virus Infectivity Titration” section.

#### p14-mediated syncytia formation on viral resistance to neutralizing antibody

Vero cells were seeded in 24-well plates and transfected the following day with the p14-expressing plasmid under a Tet-On inducible system. After 24 hours, cells were inoculated with either reporter VSV or influenza A virus (PR8) at MOI of 0.002 and 0.02, respectively, for 1 hour. Following viral adsorption, the inoculum was removed and replaced with fresh medium containing doxycycline (Dox, 0.5 µg/mL) to induce p14 expression and the appropriate neutralizing mAb. Cells were incubated for approximately 20 hours post-infection. After incubation, cells were fixed with 10% NBF and imaged using a high-content imaging system to assess syncytia formation and viral infection. Monoclonal antibodies used in this assay were either purified in-house or obtained from commercial sources.

#### Sequence Alignment and phylogenetic tree

S sequence alignment and phylogenetic tree were generated by using full-length S protein sequence with Clustal Omega. Sequences were acquired from NCBI with accession numbers: OC43-CoV, AXX83381.1; MERS-CoV, YP_009047204.1; SARS-CoV, YP_009825051.1; SARS-CoV-2, YP_009724390.1; Bat-RatG13, QHR63300.2.

#### Biosafety statement

We generated and used recombinant SARS-CoV-2 in accordance with NIH biosafety policies and with approval by the National Institutes of Health (NIH) Institutional Biosafety Committee (IBC; protocol RD-22-XI-11) and the NIH Dual Use Research of Concern Institutional Review Entity (DURC-IRE). All experiments involving infectious recombinant SARS-CoV-2 were performed in certified Biosafety Level 3 (BSL-3) laboratories by personnel trained in BSL-3 practices and procedures. Animal studies involving recombinant SARS-CoV-2 infection were conducted under protocols approved by the U.S. Food and Drug Administration White Oak and the National Institutes of Health Animal Program Animal Care and Use Committee (Animal Study Proposal: LVD-5E, LVD-11E) and performed in ABSL-3 containment.

The recombinant viruses used in this study were generated to investigate the role of spike protein-mediated syncytia formation in viral fitness, interferon resistance, and immune evasion. The work did not involve enhancing, host range, or pathogenicity beyond that required to address the scientific objectives. The anticipated benefits of this research include improved understanding of mechanisms driving SARS-CoV-2 evolution and immune escape, which informs surveillance efforts and preparedness for future emerging viral pathogens. These scientific and public health benefits were determined by the relevant biosafety committees to outweigh the potential risks when conducted under the approved biosafety and biosecurity oversight framework.

#### Statistical analysis

All statistical tests were performed as described in the figure legends using Prism v9(GraphPad Software, Inc.). Statistical significance is set as P < 0.05, and P values are indicated with: NS, not significant; *P < 0.05; **P < 0.01; ***P < 0.001, ****P < 0.0001.

## Data availability

All data supporting the conclusions of this study can be found in the article, supplementary, and source data files. Source data are provided in this paper.

## Code Availability

Custom code used for the analysis of data is available on GitHub: https://github.com/fdlts/SARS-CoV-2-Syncytia-Project

## ACKNOWLEDGMENTS

We thank Zene Matsuda at the Institute of Medical Science (University of Tokyo) for providing dual split protein system: DSP _1-7_ and DSP _8-11_, and Chhing-Wen Chang (University of Massachusetts Chan Medical School) for providing A549plusC3 cell line, Sarah Anzick (RTB, NIAID) for NSG support, Bernard Lafont, Johnson Reed, and Nicole Lackemeyer (NIAID SARS-CoV-2 Virology Core BSL-3 facility), for superb support, training, and assistance. The contributions of the NIH authors were made as part of their official duties as NIH federal employees, are in compliance with agency policy requirements, and are considered works of the U.S. government. However, the findings and conclusions presented in this paper are those of the authors and do not necessarily reflect the views of the NIH or the U.S. Department of Health and Human Services.

## Author contributions

T.L. conceived the study and designed and performed the experiments. T.L., I.K.(Insung Kang), M.K., Z.Y., and H.X. performed in vivo experiments. J.Y. performed IHC staining. K.T. assessed the pathology score. T.L., Z.H., G.S., analyzed syncytia formation in the infection of cultured epithelial cells in ALI and of mice. T.L. and J.H. generated stable cell lines. J.G. performed the analysis of SARS-CoV-2 variants proportion. I.K.(Ivan Kosik), A.C. and R.F.J. provided essential resource and scientific input. C.Y. and L.M-S. provided BAC reverse genetic system for recombinant SARS-CoV-2 generation. T.L. drafted the manuscript. T.L. and J.W.Y. reviewed and revised the manuscript. J.W.Y. supervised the study and secured funding from the Intramural Research Program of the National Institute of Allergy and Infectious Diseases (NIAID), National Institutes of Health (NIH). All the authors reviewed and proofread the manuscript.

