## Supplementary Data for "Viral Syncytia Evolve to Resist Interferon"

#### Suppl. Fig. 1

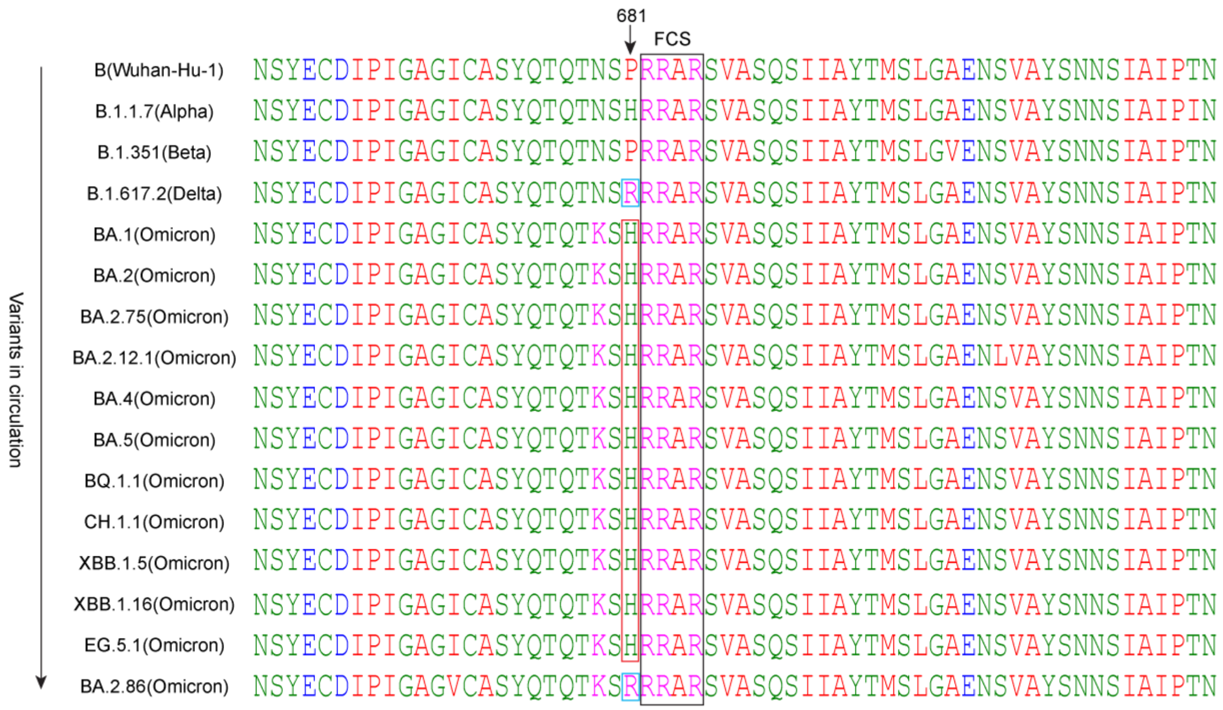
681
FCS
Variants in circulation
B(Wuhan-Hu-1) NSYECDIPIGAGICASYQTQTNSPRRARSVASQSIIAYTMSLGAENSVAYSNNSIAIPTN
B.1.1.7(Alpha) NSYECDIPIGAGICASYQTQTNSHRRARSVASQSIIAYTMSLGAENSVAYSNNSIAIPIN
B.1.351(Beta) NSYECDIPIGAGICASYQTQTNSPRRARSVASQSIIAYTMSLGVENSVAYSNNSIAIPTN
B.1.617.2(Delta) NSYECDIPIGAGICASYQTQTNSRRRARSVASQSIIAYTMSLGAENSVAYSNNSIAIPTN
BA.1(Omicron) NSYECDIPIGAGICASYQTQTKSHRRARSVASQSIIAYTMSLGAENSVAYSNNSIAIPTN
BA.2(Omicron) NSYECDIPIGAGICASYQTQTKSHRRARSVASQSIIAYTMSLGAENSVAYSNNSIAIPTN
BA.2.75(Omicron) NSYECDIPIGAGICASYQTQTKSHRRARSVASQSIIAYTMSLGAENSVAYSNNSIAIPTN
BA.2.12.1(Omicron) NSYECDIPIGAGICASYQTQTKSHRRARSVASQSIIAYTMSLGAENLVAYSNNSIAIPTN
BA.4(Omicron) NSYECDIPIGAGICASYQTQTKSHRRARSVASQSIIAYTMSLGAENSVAYSNNSIAIPTN
BA.5(Omicron) NSYECDIPIGAGICASYQTQTKSHRRARSVASQSIIAYTMSLGAENSVAYSNNSIAIPTN
BQ.1.1(Omicron) NSYECDIPIGAGICASYQTQTKSHRRARSVASQSIIAYTMSLGAENSVAYSNNSIAIPTN
CH.1.1(Omicron) NSYECDIPIGAGICASYQTQTKSHRRARSVASQSIIAYTMSLGAENSVAYSNNSIAIPTN
XBB.1.5(Omicron) NSYECDIPIGAGICASYQTQTKSHRRARSVASQSIIAYTMSLGAENSVAYSNNSIAIPTN
XBB.1.16(Omicron) NSYECDIPIGAGICASYQTQTKSHRRARSVASQSIIAYTMSLGAENSVAYSNNSIAIPTN
EG.5.1(Omicron) NSYECDIPIGAGICASYQTQTKSHRRARSVASQSIIAYTMSLGAENSVAYSNNSIAIPTN
BA.2.86(Omicron) NSYECDIPIGAGVCASYQTQTKSRRRARSVASQSIIAYTMSLGAENSVAYSNNSIAIPTN

**Supplementary Fig. 1 Furin cleavage site(FCS) motif in SARS-CoV-2 dominant circulating variants**.

Alignment of the region of *S* gene containing FCS motif from SARS-CoV-2 dominant circulating variants in an order of emergency in time. The RXXR furin cleavage motif (682-685 in Wuhan-Hu-1) is marked in a black rectangle. The proximal residue at 681 which affects the cleavage efficiency and syncytia is indicated in arrowhead. The hallmark mutation at 681 in Delta and the new emerging Omicron subvariant BA.2.86 are marked in blue; other Omicron subvariants except BA.2.86 are marked in red. Sequences were acquired from NCBI with access number being: YP_009724390.1, QQH18533.1, WLL57934.1, UVN17823.1, UFT00449.1, UIG03312.1, USV68346.1, WMV91511.1, WKR02063.1, UPU09668.1, UWM38596.1, UYH63216.1, BES80299.1, BES80198.1, WGP26425.1, WLW39834.1 . Alignment was generated by ClustalOmega using full-length S protein sequence.

### Suppl. Fig. 2

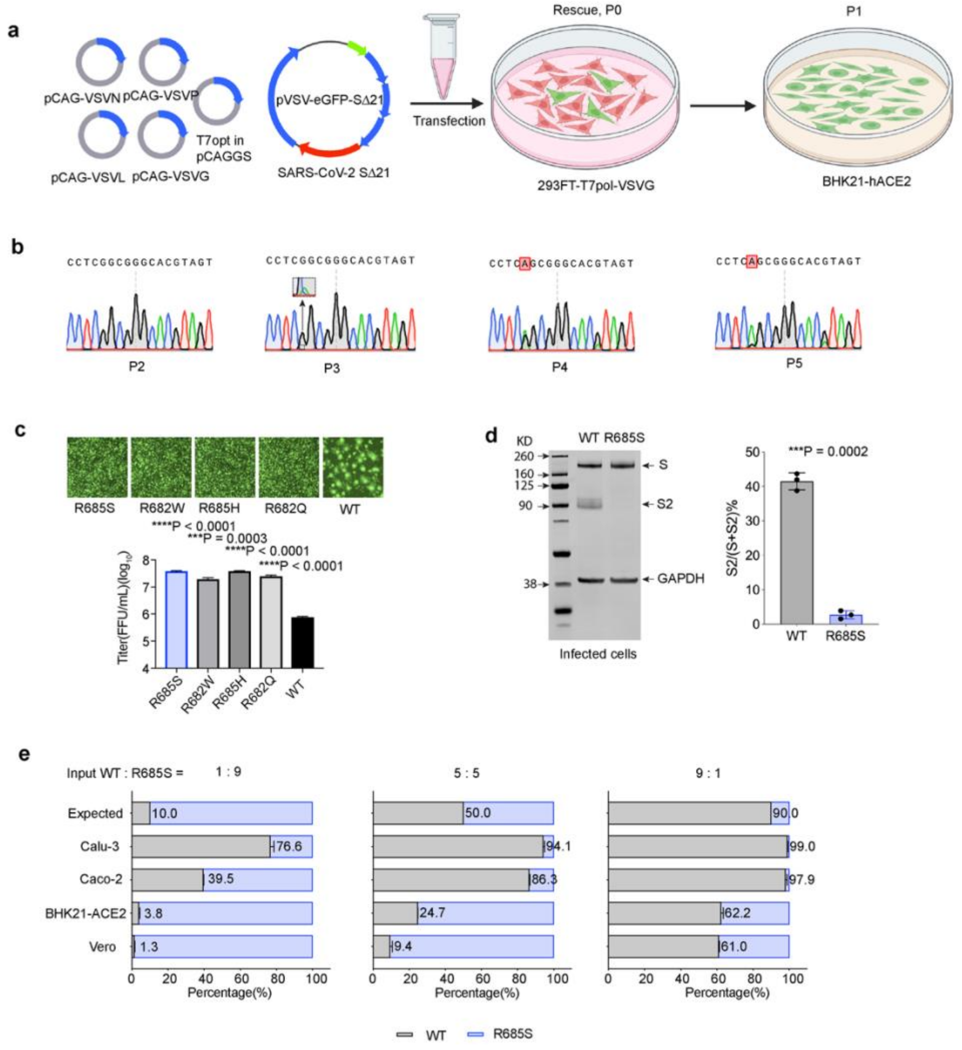
a
pCAG-VSVN
pCAG-VSVP
pCAG-VSVL
pCAG-VSVG
T7opt in pCAGGS
pVSV-eGFP-SΔ21
SARS-CoV-2 SΔ21
Transfection
Rescue, P0
293FT-T7pol-VSVG
P1
BHK21-hACE2
b
CCTCGGCGGGCACGTAGT
P2
P3
P4
P5
c
R685S
R682W
R685H
R682Q
WT
****P < 0.0001
***P = 0.0003
****P < 0.0001
****P < 0.0001
Titer(FFU/mL)(log10)
d
KD
WT R685S
S
S2
GAPDH
Infected cells
***P = 0.0002
S2/(S+S2)%
e
Input WT : R685S =
1 : 9
5 : 5
9 : 1
Expected
Calu-3
Caco-2
BHK21-ACE2
Vero
Percentage(%)
WT
R685S

**Supplementary Fig. 2 Generation of replication-competent chimeric SARS-CoV-2 S bearing VSV virus and forward genetic selection of fitness-advantaged mutants**.

**a** Stable cell line 293FT cells expressing VSVG and T7 polymerase were transfected with plasmids VSV-N, -P, -L, -G, and codon-optimized T7 polymerase and an infectious molecular cDNA of pVSV-eGFP-SARS-CoV-2-S to produce replication-competent VSV-eGFP-SARS-CoV-2-S (referred as rVSV-S), the rescued viral stock (P0) was then passaged 4 times in BHK21 cells stably expressing human ACE2 (BHK21-ACE2). Schematic diagram was created in BioRender (Created in BioRender. Gibbs, J. (2026) https://BioRender.com/81pa3si).

**b** FCS mutations were identified by Sanger sequencing on passaged rVSV- S on the whole *S* gene. The region of FCS was shown to show double peaks at G2045 (R682 at amino acid level) in *S* gene in passages 3 to 5, but not in passage 2. Another mutation at G2054(R685 at amino level) was observed in P4 and P5.

**c** Syncytia formation and titer of plaque-purified rVSV-S *wt* and mutants. Four mutants with mutations in FCS were identified in a plaque-purified assay. Representative images show decreased syncytia formation of all mutants in BHK21-ACE2 cells (top). Titer was measured in BHK21-ACE2 cells (bottom), and all FCS mutants gained growth advantage in BHK21-ACE2 cells(n=3 biological replicates). Data show mean ± s.d.

**d** Lysates of Vero cells infected with rVSV-S (wild-type or R685S mutant) were analyzed by western blotting at 24 h.p.i. The R685S mutation almost completely abolished cleavage at the FCS.

**e** Competition assay between rVSV- S *wt* and R685S at total MOI of 0.01, showing infected cells by eGFP(R685S) and mCherry (*wt*) channels. The input ratio was for inoculation, and the output infection percentage was detected by flow cytometry at 7 h.p.i (n=3 biological replicates). Data show mean ± s.d.

Statistical analysis was performed using a two-tailed, unpaired t-test. ***P < 0.001.

### Suppl. Fig. 3

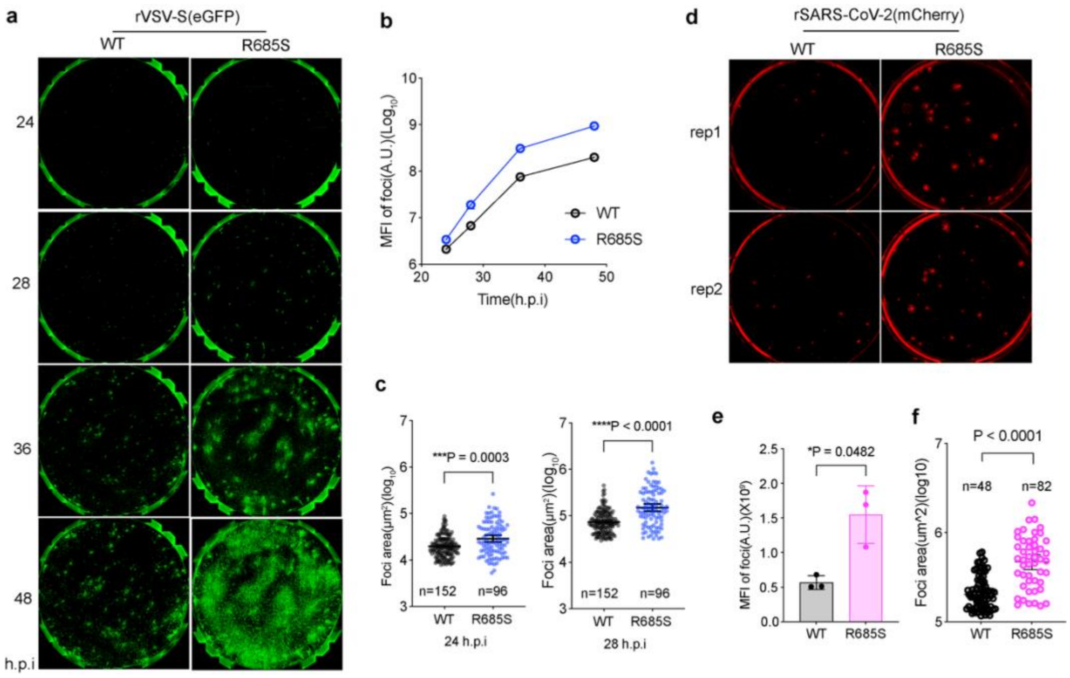

a
rVSV-S(eGFP)
WT
R685S
24
28
36
48
h.p.i
b
MFI of foci(A.U.)(Log10)
Time(h.p.i)
WT
R685S
c
Foci area(μm²)(log10)
***P = 0.0003
n=152
n=96
WT
R685S
24 h.p.i
****P < 0.0001
n=152
n=96
WT
R685S
28 h.p.i
d
rSARS-CoV-2(mCherry)
WT
R685S
rep1
rep2
e
MFI of foci(A.U.)(X10⁶)
*P = 0.0482
WT
R685S
f
Foci area(um^2)(log10)
P < 0.0001
n=48
n=82
WT
R685S

**Supplementary Fig. 3 FCS mutant R685S gains fitness advantage in Vero cell.**

**a** Time-lapse image of rVSV-S *wt* and R685S infections in Vero cell, with 100 FFU/well in a 12-well plate in inoculation.

**b** and **c** Replication dynamics of rVSV-S *wt* and R685S infections in Vero cells. Mean fluorescence intensity (MFI) of each infectious focus was measured(**b**). Data show mean ± s.d. (**b**, n=4 biological replicates). Focus area was compared between rVSV-S *wt* and R685S at 24 h.p.i(left) and 28 h.p.i(right)(**c**), n= 152 foci and 96 foci for *wt* and R685S, respectively. Data show the geometric mean ± 95% CI (**c**).

**d** Infection of recombinant rSARS-CoV-2 *wt* (WA-1 strain) and R685S mutant, with expression of mCherry gene linked to nucleoprotein via a 2A linker, in Vero cells at 36 h.p.i in a 12-well plate. Two representative images for each virus (rep1 and rep2) were shown. Cells were overlayed with 1.25% avicel after infection.

**e** Mean fluorescence intensity of infection foci as shown in **d** (n=3 biological replicates). Data show mean ± s.d.

**f** Individual area of infection foci as shown in d, n= 48 and 82 foci for *wt* and R685S, respectively. Data show the geometric mean ± 95% CI. Each dot represents one focus in **c** and **f**. For **c** and **f**, statistical analysis was performed using a two-sided, unpaired Mann-Whitney test. *P < 0.05, **P < 0.01, ***P < 0.001 and ****P < 0.0001; NS, not significant.

#### Suppl. Fig. 4

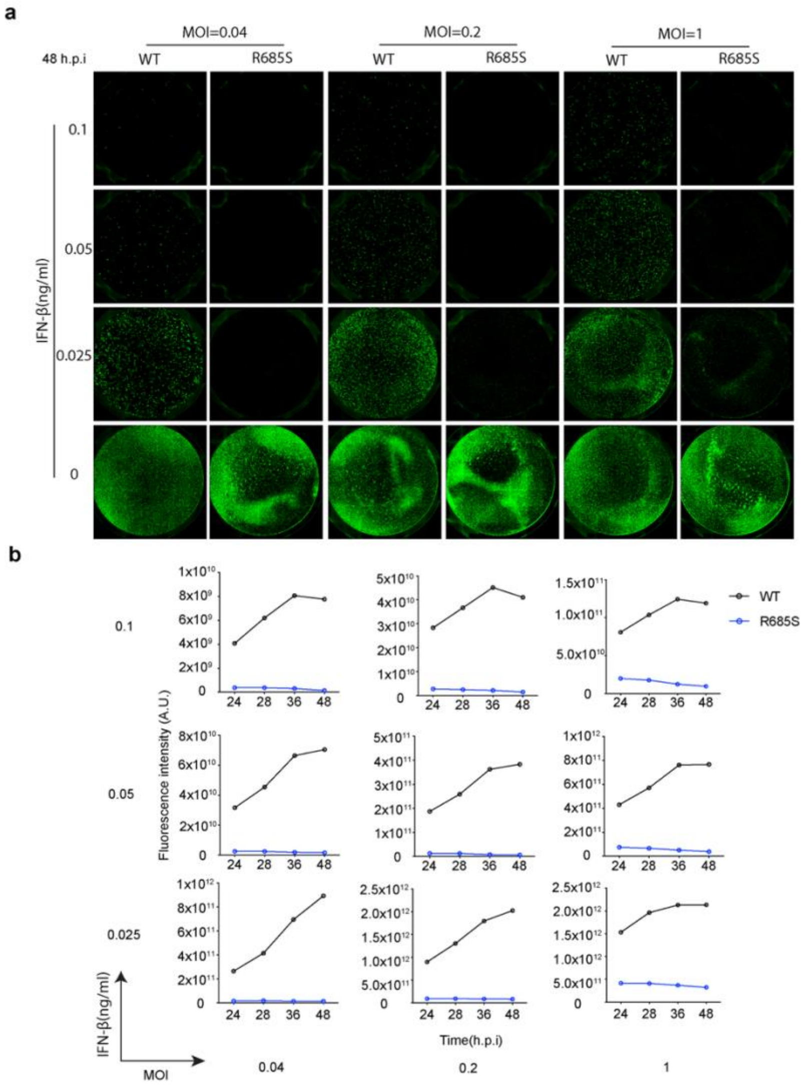
a
MOI=0.04
MOI=0.2
MOI=1
48 h.p.i
WT
R685S
IFN-β(ng/ml)
0.1
0.05
0.025
0
b
WT
R685S
Fluorescence intensity (A.U.)
Time(h.p.i)
24 28 36 48
IFN-β(ng/ml)
MOI
0.04
0.2
1

**Supplementary Fig. 4 Sensitivity of *wt* and R685S to IFN-β in Vero cells in VSV system.**

**a** Representative images of rVSV-S *wt* and R685S infections under exogenous IFN-β treatment in Vero cells at 48 h.p.i, in 12-well plates. Cells were pretreated with IFN-β for 18-20 h prior to infection, and corresponding IFN was maintained throughout the infection.

**b** Fluorescence intensity of each infection with different MOI and various concentrations of IFN-β.

**Suppl. Fig. 5**

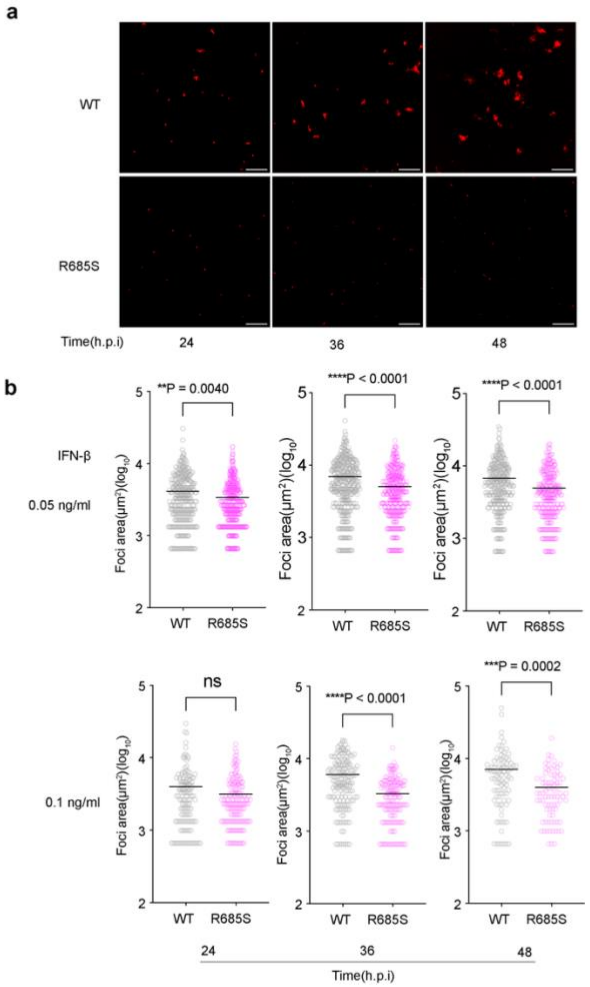
a
WT
R685S
Time(h.p.i)
24
36
48
b
**P = 0.0040
****P < 0.0001
****P < 0.0001
IFN-β
0.05 ng/ml
Foci area(μm²)(log10)
WT
R685S
ns
****P < 0.0001
***P = 0.0002
0.1 ng/ml
Time(h.p.i)

**Supplementary Fig. 5 Sensitivity of *wt* and R685S to IFN-β in Vero cells in SARS-CoV-2 system.**

**a** Representative images of rSARS-CoV-2 *wt* and R685S infections under exogenous IFN-β (0.1 ng/ml) treatment in Vero cells at MOI of 0.01. Cells were pretreated with IFN-β for 18-20 h prior to infection, and corresponding IFN was maintained throughout the infection. Scale bar, 500 μm.

**b** Infection focus area of rSARS-CoV-2 *wt* and R685S under IFN-β treatment at the concentration of 0.05 (up) and 0.1 (bottom) ng/ml. Data show mean ± s.e.m. each dot represents one infection focus. Statistical analysis was performed using a two-sided, unpaired Mann-Whitney test. ***P < 0.001 and ****P < 0.0001; ns, not significant.

### Suppl. Fig. 6

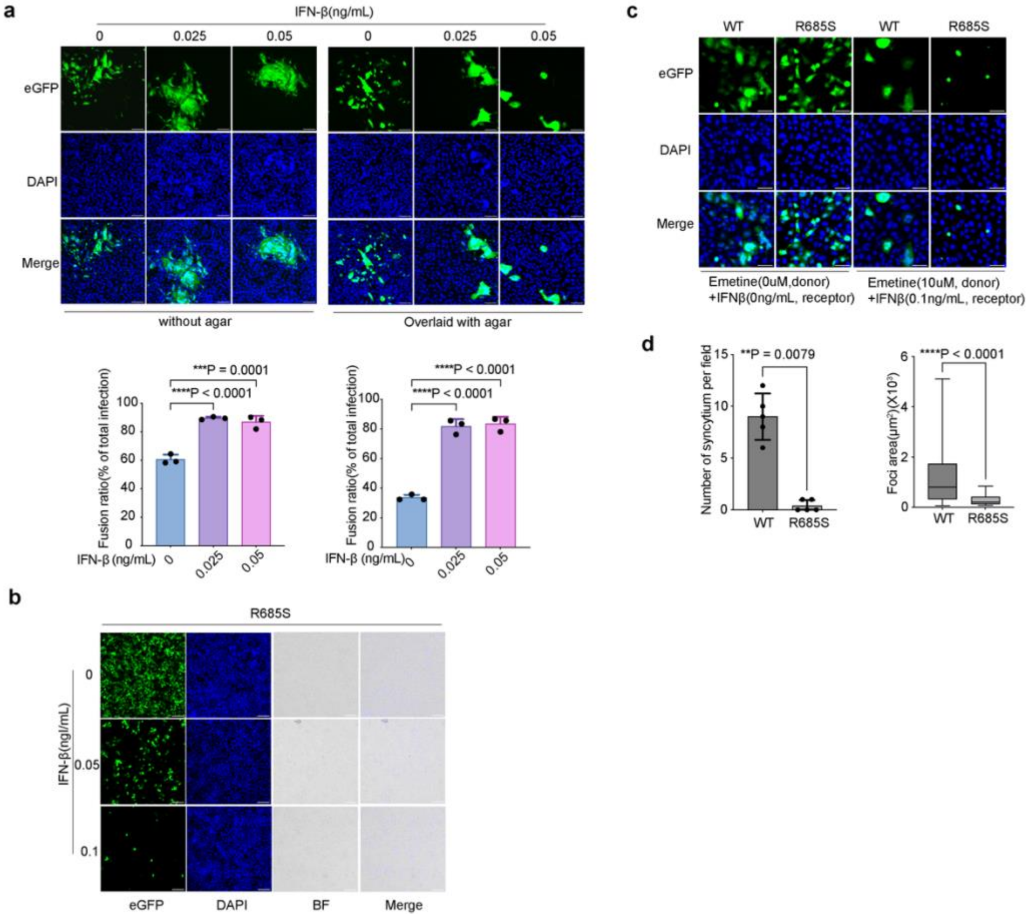
a
IFN-β(ng/mL)
0
0.025
0.05
0
0.025
0.05
eGFP
DAPI
Merge
without agar
Overlaid with agar
***P = 0.0001
****P < 0.0001
Fusion ratio(% of total infection)
IFN-β (ng/mL)
****P < 0.0001
****P < 0.0001
b
R685S
IFN-β(ngl/mL)
0
0.05
0.1
eGFP
DAPI
BF
Merge
c
WT
R685S
WT
R685S
eGFP
DAPI
Merge
Emetine(0uM,donor) +IFNβ(0ng/mL, receptor)
Emetine(10uM, donor) +IFNβ(0.1ng/mL, receptor)
d
**P = 0.0079
Number of syncytium per field
WT
R685S
****P < 0.0001
Foci area(μm²)(X10³)
WT
R685S

**Supplementary Fig. 6 Syncytia but not cell-free spread confers virus resistance to IFN-β anti-viral activity.**

**a** Effects of IFN-β on rVSV-S *wt* fusion ratio in Vero cells. Pretreated Vero cells by IFN-β were infected by rVSV-S, cells were overlaid without(left) or with(right) agar for 24 h with IFN-β in the medium. Infected cells were fixed and scanned by the high-content imaging system Cytation 5. Fusion ratio (%) was calculated by dividing the infection area of syncytia by the total infected area. In Gen 5: the size of object was set to >60 μm to calculate the area of syncytia, which is about 3-fold the size of a single Vero cell *in situ* (n=3 biological replicates). Data show mean ± s.d.

**b** Representative images of rVSV- S R685S infection in Vero cells under IFN-b, linked to images in Fig.2h for comparison. BF means bright field, and Merge was conducted between BF and DAPI to show syncytia formation.

**Suppl. Fig. 7**

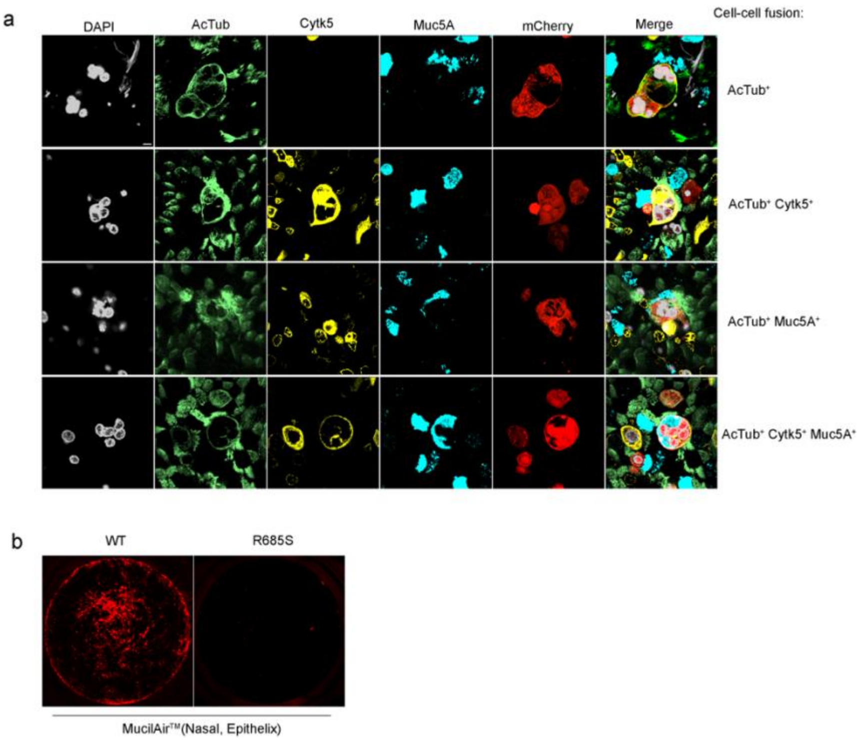
a
DAPI
AcTub
Cytk5
Muc5A
mCherry
Merge
Cell-cell fusion:
AcTub+
AcTub+ Cytk5+
AcTub+ Muc5A+
AcTub+ Cytk5+ Muc5A+
b
WT
R685S
MucilAir™(Nasal, Epithelix)

**Supplementary Fig. 7 SARS-CoV-2 infection in human respiratory tract epithelial cells in air-liquid interface format**.

**a** Cell types in syncytia formation, related to Fig. 3g. Infected small airway ALI were stained with anti-AcTub (green signal, marker for ciliated cell), anti-Cytk5 (yellow signal, marker for basal cell) and anti-Muc5A (cyan signal, marker for goblet cell) Abs and counterstained with DAPI (gray signal). Individual syncytia were shown from 3D reconstructed confocal images. Scale bar, 5μm.

b Entire Nasal epithelial cell ALI infected by rSARS-CoV-2 *wt* (left) and R685S mutant(right), related to Fig. 3h.

### Suppl. Fig. 8

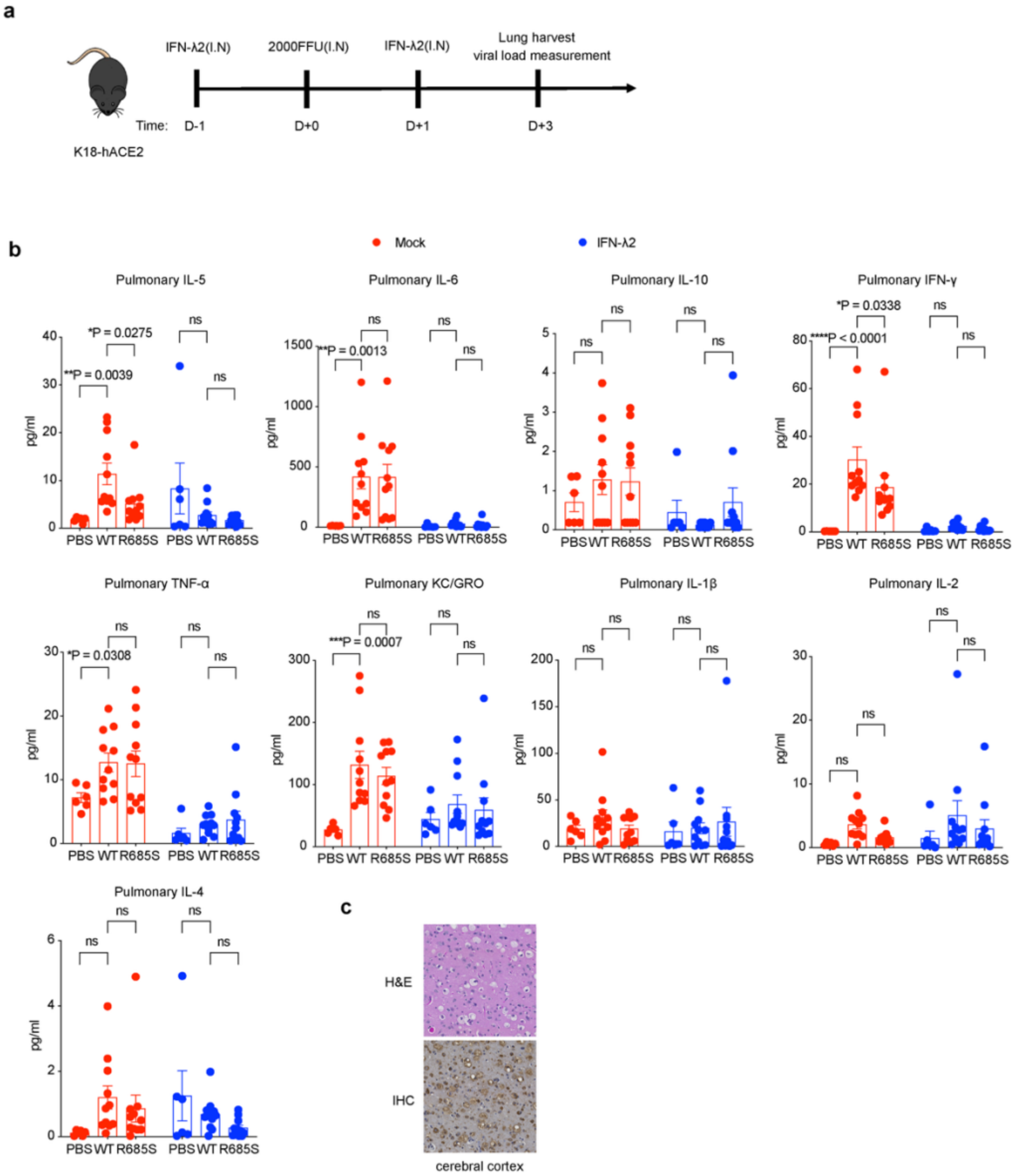

a
IFN-λ2(I.N)
2000FFU(I.N)
IFN-λ2(I.N)
Lung harvest
viral load measurement
Time:
D-1
D+0
D+1
D+3
K18-hACE2
b
Mock
IFN-λ2
Pulmonary IL-5
Pulmonary IL-6
Pulmonary IL-10
Pulmonary IFN-γ
Pulmonary TNF-α
Pulmonary KC/GRO
Pulmonary IL-1β
Pulmonary IL-2
Pulmonary IL-4
pg/ml
PBS WT R685S PBS WT R685S
**P = 0.0039
*P = 0.0275
**P = 0.0013
*P = 0.0338
****P < 0.0001
*P = 0.0308
***P = 0.0007
ns
c
H&E
IHC
cerebral cortex

**Supplementary Fig. 8 Proinflammation cytokine profile in lung tissue of K18-hACE2 mice infected by rSARS-CoV-2 *wt* and R685S mutant**.

**a** Schematic workflow (created by Adobe Illustrator) of rSARS-CoV-2 infection in K18 mice treated with murine IFN-λ2. I.N, Intranasal.

**b** Proinflammation cytokine expression level in lung tissues of infected K18 mice with or without IFN-λ2 treatment. Data show the combination of two independent experiments. Data show mean ± s.d (n=11 mice).

**c** Representative images of HE and IHC staining of R685S infection in cerebral cortex of the brain tissue.

Statistical analysis was performed using two-way ANOVA (**b**). *P < 0.05, **P < 0.01, ***P < 0.001 and ****P < 0.0001; ns, not significant.

#### Suppl. Fig. 9

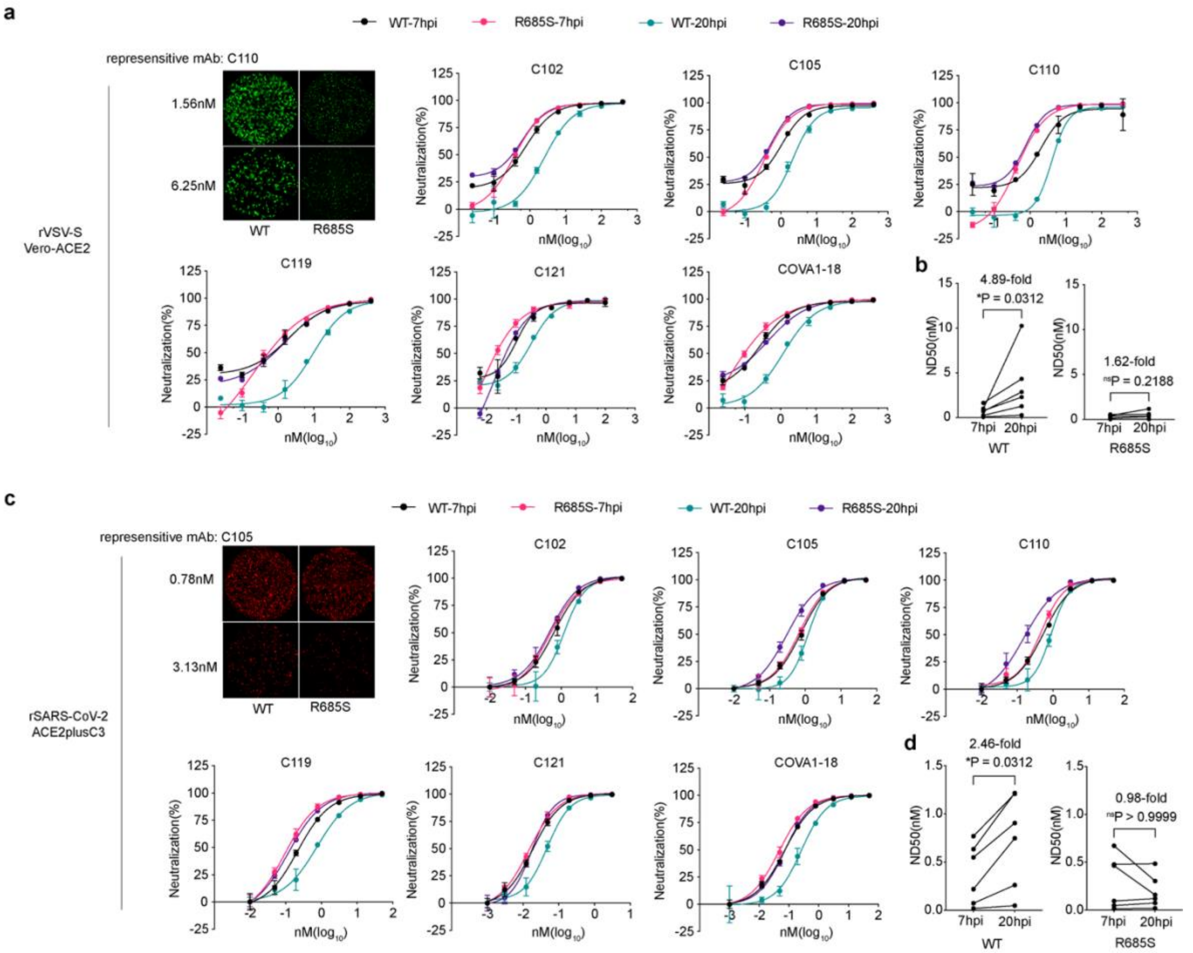
a
WT-7hpi
R685S-7hpi
WT-20hpi
R685S-20hpi
represensitive mAb: C110
1.56nM
6.25nM
WT
R685S
rVSV-S
Vero-ACE2
C102
C105
C110
C119
C121
COVA1-18
Neutralization(%)
nM(log10)
b
ND50(nM)
4.89-fold
*P = 0.0312
7hpi
20hpi
WT
1.62-fold
nsP = 0.2188
R685S
c
represensitive mAb: C105
0.78nM
3.13nM
rSARS-CoV-2
ACE2plusC3
d
2.46-fold
*P = 0.0312
0.98-fold
nsP > 0.9999

**Supplementary Fig. 9 Monoclonal antibody neutralization profile against *wt* and R685S mutant infection**.

A two-timepoints assay was performed to assess the contribution of syncytia spread of virus to the neutralization effect.

**a** and **b** Neutralization profile of 6 potent mAbs targeting RBD of S against rVSV-S *wt* and R685S infection in Vero-ACE2 cells. Representative images of rVSV-S *wt* and R685S infection at 20 h.p.i were shown (**a**, top left panel). Raw neutralization data were performed in nonlinear regression analysis (**a**, top right and bottom panel). 50% neutralization dose ($ND_{50}$) of each antibody was determined and shown (**b**).

**c** and **d** Neutralization profile of 6 potent mAbs against rSARS2 *wt* and R685S infection in ACE2plusC3 cells. Representative images of rSARS-CoV-2 *wt* and R685S infection at 20 h.p.i were shown (**c**, top left panel). The neutralization curve was fitted with nonlinear regression analysis, and the $ND_{50}$ of each antibody was determined(**d**).

Data show mean ± s.d (n=3 wells per dose). Statistical analysis was performed using a two-sided Wilcoxon matched pair signed rank test. *P < 0.05; ns, not significant.

**Suppl. Fig. 10**

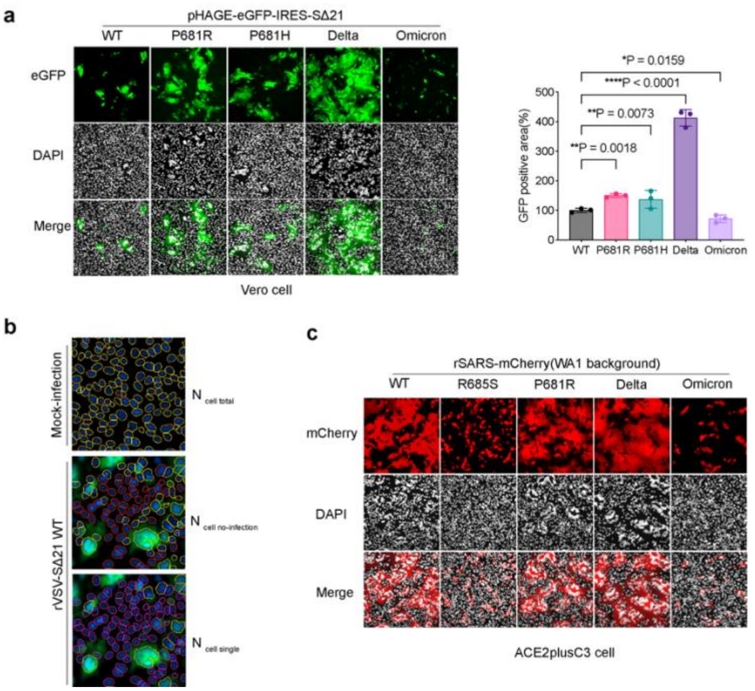
a
pHAGE-eGFP-IRES-SΔ21
WT
P681R
P681H
Delta
Omicron
eGFP
DAPI
Merge
Vero cell
*P = 0.0159
****P < 0.0001
**P = 0.0073
**P = 0.0018
GFP positive area(%)
500
400
300
200
100
0
WT
P681R
P681H
Delta
Omicron
b
Mock-infection
rVSV-SΔ21 WT
N cell total
N cell no-infection
N cell single
c
rSARS-mCherry(WA1 background)
WT
R685S
P681R
Delta
Omicron
mCherry
DAPI
Merge
ACE2plusC3 cell

**Supplementary Fig. 10 Syncytia formation mediated by S variants.**

**a** Syncytia formation in Vero cells mediated by S-expressing plasmids, containing *wt*, P681R, P681H, Delta or Omicron (BA.1) S. Representative images at 20 h.p.t (left) and GFP positive area normalized to *wt* (right) was shown. Data show mean ± s.d (n=3 biological replicates). Scale bar, 100 μm.

**b** fusion index calculation workflow, fusion index was defined as described in the Method section. Scale bar, 100 μm.

**c** Syncytia formation in ACE2plusC3 cells mediated by rSARS-CoV-2 virus infection, containing *wt*, R685S, P681R, Delta or Omicron (BA.1) S. Representative images at 20 h.p.i were shown. Scale bar, 100 μm. Statistical analysis was performed using one-way ANOVA. *P < 0.05, **P < 0.01, ***P < 0.001 and ****P < 0.0001; ns, not significant.

#### Suppl. Fig. 11

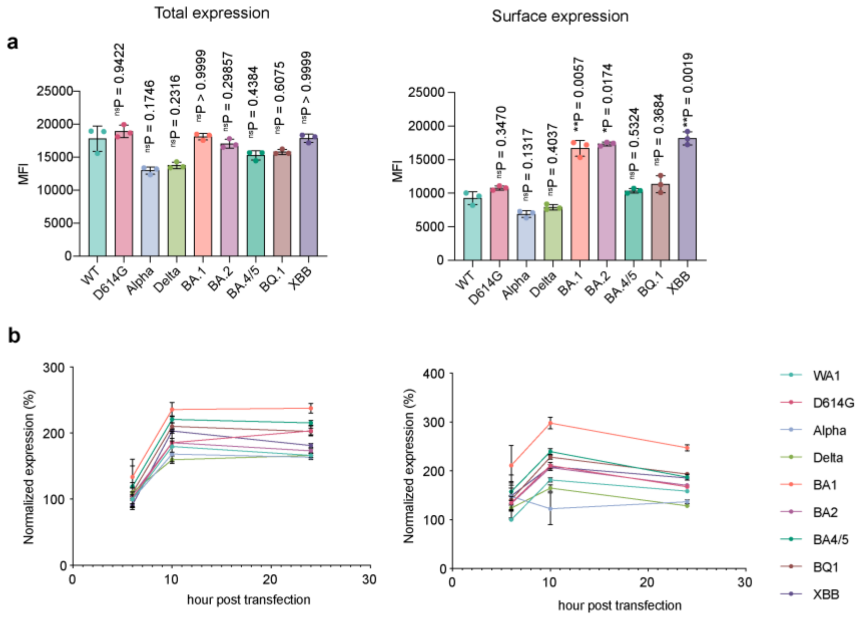
a
Total expression
Surface expression
MFI
nsP = 0.9422
nsP = 0.1746
nsP = 0.2316
nsP > 0.9999
nsP = 0.29857
nsP = 0.4384
nsP = 0.6075
nsP > 0.9999
nsP = 0.3470
nsP = 0.1317
nsP = 0.4037
**P = 0.0057
*P = 0.0174
nsP = 0.5324
nsP = 0.3684
**P = 0.0019
WT
D614G
Alpha
Delta
BA.1
BA.2
BA.4/5
BQ.1
XBB
b
Normalized expression (%)
hour post transfection
WA1
D614G
Alpha
Delta
BA1
BA2
BA4/5
BQ1
XBB

**Supplementary Fig. 11 Protein expression level of S variants in Vero and 293FT cells**

**a** S expression level of variants in Vero cells. S-containing plasmid were transfected to Vero cells. At 10 h.p.i., cells were detached and fixed with 4%PFA. Total S expression (left, permeabilized by 0.2% saponin) and cell surface S expression (right, without permeabilization) were determined by flow cytometry stained by antibody S2P6 targeting stem region of S, at 1μg/ml. Mean fluorescence intensity (MFI) of positive cells was shown.

**b** S expression kinetics of variants in 293FT cells. As descripted in **a**, S expression level in 293FT cells was determined at indicated time point post transfection.

**Suppl. Fig. 12**

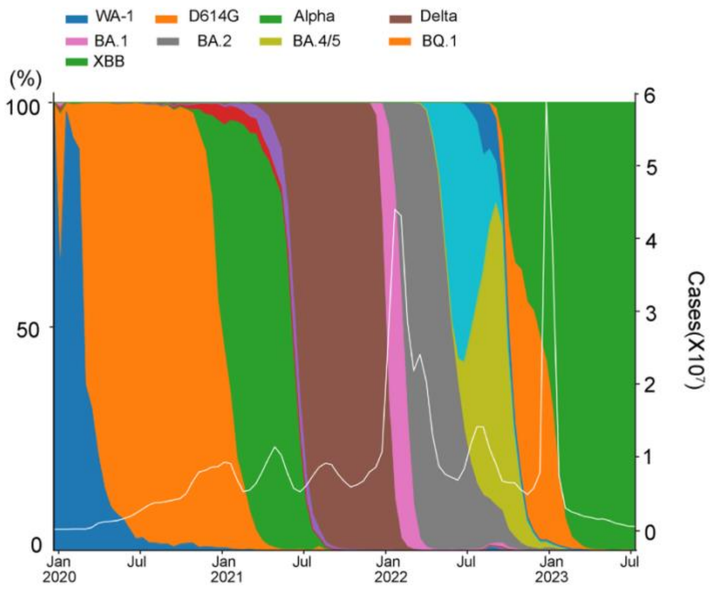


**Supplementary Fig. 12 Proportion of dominant SARS-CoV-2 variants over time**.

Daily global reported infection case and vaccination datasets were downloaded from Our World in Data (https://ourworldindata.org/covid-vaccinations) as of July 2023. The dataset was plotted by Python program. The white line shows daily new confirmed COVID-19 cases in 7-day rolling average.

**Suppl. Fig. 13**

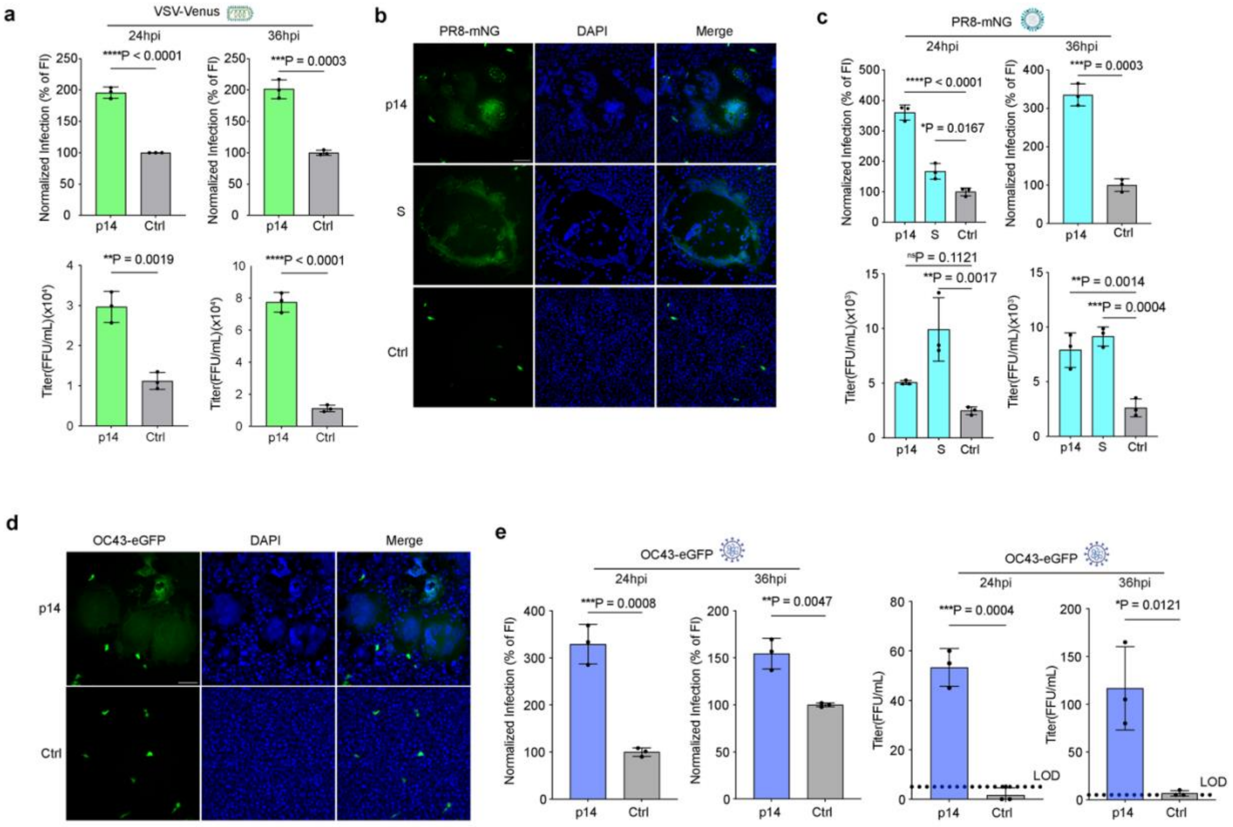
a
VSV-Venus
24hpi
36hpi
****P < 0.0001
***P = 0.0003
Normalized Infection (% of FI)
**P = 0.0019
****P < 0.0001
Titer(FFU/mL)(x10⁴)
p14
Ctrl
b
PR8-mNG
DAPI
Merge
p14
S
Ctrl
c
PR8-mNG
24hpi
36hpi
****P < 0.0001
*P = 0.0167
***P = 0.0003
nsP = 0.1121
**P = 0.0017
**P = 0.0014
***P = 0.0004
Titer(FFU/mL)(x10³)
d
OC43-eGFP
DAPI
Merge
p14
Ctrl
e
OC43-eGFP
24hpi
36hpi
***P = 0.0008
**P = 0.0047
***P = 0.0004
*P = 0.0121
Titer(FFU/mL)
LOD

**Supplementary Fig. 13 p14-mediated syncytia formation enhances virus resistance to IFN**

**a** Fluorescence intensity (top) and titer (bottom) of VSV infection in Vero cells with syncytia formation mediated by p14, under IFN-β 0.2 ng/ml treatment. n = 3 biological replicates.

**b** Representative images of PR8-mNG infection with p14-mediated syncytia formation in Vero cell treated with 1 ng/mL IFN-β**.** Scale bar, 100μm.

**c** Fluorescence intensity (top) and titer (bottom) of PR8-mNG infection in Vero cells with syncytia formation mediated by p14, under IFN-β 0.5 ng/ml treatment. n = 3 biological replicates.

**d** Representative images of OC43-eGFP infection with p14-mediated syncytia formation in Vero cell treated with 5 ng/mL IFN-β**.** Scale bar, 100μm.

**e** Fluorescence intensity (top) and titer (bottom) of OC43 infection in Vero cells with syncytia formation mediated by p14, under IFN-β 1 ng/ml treatment. n = 3 biological replicates.

Statistical analyses were performed using an unpaired t test (**a**, **c**(upper right panel), **e**) or one-way ANOVA with multiple comparisons (**c**, upper right panel excluded)). *P < 0.05, **P < 0.01, ***P < 0.001 and ****P < 0.0001; ns, not significant. Data show mean ± s.d.

### Suppl. Fig. 14

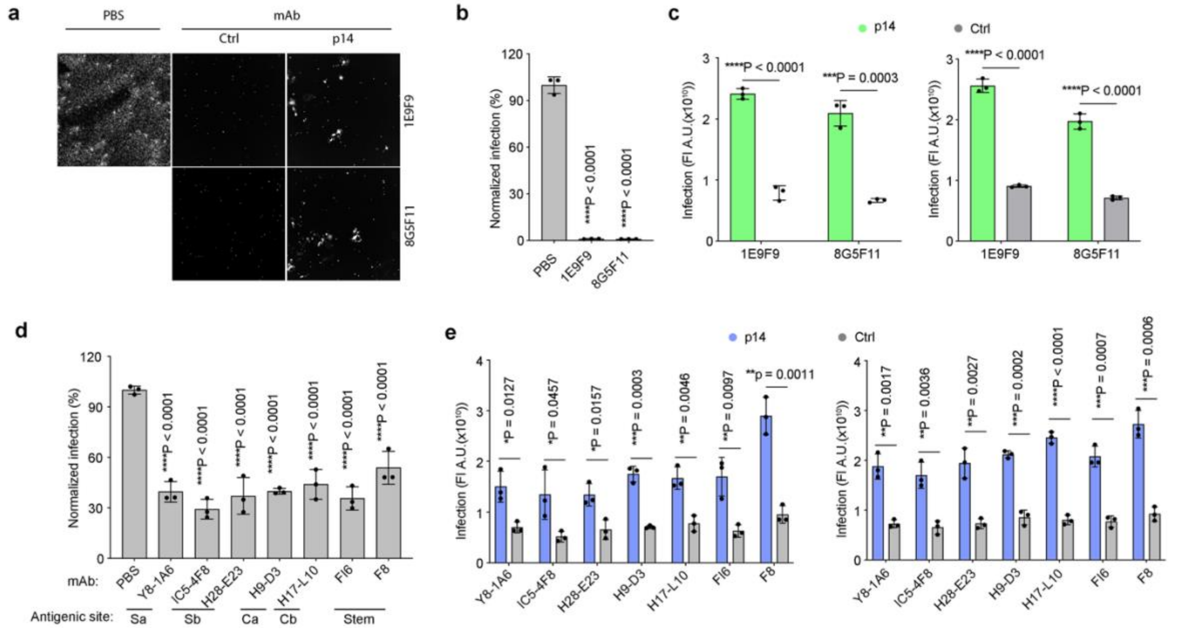

a
PBS
mAb
Ctrl
p14
1E9F9
8G5F11
b
Normalized infection (%)
PBS
1E9F9
8G5F11
****P < 0.0001
****P < 0.0001
c
p14
Ctrl
Infection (FI A.U.(x10^10))
****P < 0.0001
***P = 0.0003
****P < 0.0001
****P < 0.0001
1E9F9
8G5F11
d
Normalized infection (%)
mAb:
PBS
Y8-1A6
IC5-4F8
H28-E23
H9-D3
H17-L10
FI6
F8
Antigenic site:
Sa
Sb
Ca
Cb
Stem
e
p14
Ctrl
Infection (FI A.U.(x10^10))
*P = 0.0127
*P = 0.0457
*P = 0.0157
***P = 0.0003
**P = 0.0046
**P = 0.0097
**P = 0.0011
**P = 0.0017
**P = 0.0036
**P = 0.0027
***P = 0.0002
****P < 0.0001
***P = 0.0007
***P = 0.0006

**Supplementary Fig. 14 p14-mediated syncytia formation enhances virus resistance to neutralizing antibodies**

**a** Representative image of VSV infection with p14-mediated syncytia formation in Vero cell with neutralizing antibody 1E9F9 and 8G5F11 at 5 μg/ml at 20 h.p.i. Scale bar, 500 μm.

**b** Inhibition of antibody 1E9F9 and 8G5F11 at 5 μg/ml on VSV infection in Vero cells at 20 h.p.i. n = 3 biological replicates.

**c** Infection of VSV in Vero cells transfected with p14, treated with antibody 1E9F9 and 8G5F11 at concentration of 50 μg/ml (left) or 5 μg/ml (right) for 20 h.p.i. n = 3 biological replicates.

**d** Inhibition of monoclonal antibodies at 5 μg/ml on PR8 infection in Vero cells at 20 h.p.i. n = 3 biological replicates.

**e** Infection of PR8 in Vero cells transfected with p14, treated with monoclonal antibody at concentration of 50 μg/ml (left) or 5 μg/ml (right) for 20 h.p.i. n = 3 biological replicates.

Statistical analyses were performed using an unpaired t test (**b**, **c**, **e**) or one-way ANOVA with multiple comparisons (**d**). *$P < 0.05$, **$P < 0.01$, ***$P < 0.001$ and ****$P < 0.0001$. Data show mean ± s.d.
